# High throughput screening identifies SOX2 as a Super Pioneer Factor that inhibits DNA methylation maintenance at its binding sites

**DOI:** 10.1101/2020.02.10.941682

**Authors:** Ludovica Vanzan, Hadrien Soldati, Victor Ythier, Santosh Anand, Nicole Francis, Rabih Murr

## Abstract

Access of mammalian transcription factors (TFs) to regulatory regions, an essential event for transcription regulation, is hindered by chromatin compaction involving nucleosome wrapping, repressive histone modifications and DNA methylation. Moreover, methylation of TF binding sites (TBSs) affects TF binding affinity to these sites. Remarkably, a special class of TFs called pioneer transcription factors (PFs) can access nucleosomal DNA, leading to nucleosome remodelling and chromatin opening. However, whether PFs can bind to methylated sites and induce DNA demethylation is largely unknown.

Here, we set up a highly parallelized approach to investigate PF ability to bind methylated DNA and induce demethylation. Our results indicate that the interdependence between DNA methylation and TF binding is more complex than previously thought, even within a select group of TFs that have a strong pioneering activity; while most PFs do not induce changes in DNA methylation at their binding sites, we identified PFs that can protect DNA from methylation and PFs that can induce DNA demethylation at methylated binding sites. We called the latter “super pioneer transcription factors” (SPFs), as they are seemingly able to overcome several types of repressive epigenetic marks. Importantly, while most SPFs induce TET-dependent active DNA demethylation, SOX2 binding leads to passive demethylation by inhibition of the maintenance methyltransferase DNMT1 during replication. This important finding suggests a novel mechanism allowing TFs to interfere with the epigenetic memory during DNA replication.

## Introduction

Transcription factor (TF) binding to specific sites at gene-proximal and -distal regulatory regions is a fundamental step in gene expression regulation. However, as the sequence of a given regulatory region is similar in all cells, the regulation of cell-specific transcription is likely to depend on non-genetic regulators. Indeed, TF access to regulatory elements is controlled by the chromatin structure, which in turn is modulated by epigenetic modifications (nucleosome remodelling, histone modifications and DNA methylation). These are classified into active or repressive according to their effect on gene expression. DNA tightly wrapped around nucleosomes due to repressive histone marks was shown to be refractory to TF binding^1^. Therefore, activity of nucleosome remodellers to open the chromatin was deemed necessary prior to TF binding to compact chromatin^2^. However, several studies identified a new class of TFs, called pioneer transcription factors (PFs) that access their target sites in condensed chromatin. This results in chromatin “de-condensation” by nucleosome remodelling, recruitment of “settler” TFs that are unable to access condensed chromatin, and transcription activation^3–6^. These findings hint for a more complex relationship between epigenetic- and genetic-based mechanisms in transcription regulation than previously thought. It is therefore important to establish whether and when epigenetic mechanisms constitute a primary event in the regulation of transcription and when do they simply result from previous events governed by the genetic composition of the regulatory regions (i.e. TF binding).

DNA methylation is an essential epigenetic modification that was hypothesized not only to inhibit the accessibility of DNA but also its affinity to TFs^7, 8^. Remarkably, recent studies have shown that not all TFs show sensitivity to DNA methylation. Moreover, some TFs were shown to preferentially bind to methylated sites^9–11^. However, it is currently less clear whether TFs, notably those that can bind to methylated DNA, lead to changes in the methylation status of their binding sites (i.e. DNA demethylation), and if so, how. More precisely, can PFs, in addition to their ability to remodel the nucleosomes, induce DNA demethylation?

Using a high throughput approach, our study methodically determined the ability of reported PFs to induce DNA demethylation at their binding sites in mouse embryonic stem cells (ESCs) and *in vitro* differentiated neuronal progenitors (NPs). Results show that, while many PFs do not affect the methylation status of their binding sites, a group of PFs that we call “protective pioneer transcription factors (PPFs)” protect DNA from *de novo* acquisition of methylation, while another group called “super pioneer transcription factors (SPFs)” induce DNA demethylation at their methylated binding sites. Importantly, we show that most SPF-driven demethylation is TET-dependent, except for SOX2 that inhibits DNMT1 at the replication fork, thus leading to replication-dependent passive DNA demethylation.

## Results

### Hi-TransMet: a high throughput assay for the analysis of transcription factor binding effect on DNA methylation

We developed a method called Hi-TransMet (for <u>hi</u>gh throughput analysis of <u>trans</u>cription factor effect on DNA <u>met</u>hylation) that allows to assess the effect of up to hundreds of transcription factors (TFs) on DNA methylation around their binding sites, in parallel. The method is based on bisulfite deep sequencing of PCR amplicons of the same sequence backbone with known methylation status but containing different TF binding motifs. Specifically, we used a transgenic mouse ESC line, in which a targeting site surrounded by two inverted LoxP sites, was engineered at the β-globin locus^12–14^. This locus is inactive in non-erythroid cells and was shown to not interfere with the methylation status of the cassette. Using the Recombinase-Mediated Cassette Exchange (RMCE) approach^15^, we could replace the cassette with any DNA fragment surrounded by two LoxP sites in a donor plasmid (**supplementary figure 1a**). In particular, we chose as a backbone a bacterial DNA fragment (FR1, **supplementary figure 1b**) with a CpG ratio of 3.6%, making it a CpG island akin to an intermediate CpG-content promoter (ICP)^16^. FR1 was reported to get fully methylated, when inserted in the RMCE site either unmethylated or *in vitro* methylated (M.SssI methyltransferase treatment). Importantly, the fragment is unlikely to contain any mammalian TF motif due to evolutionary distance to mammals^12^, thus inclusion of a TF binding motif within it could address whether the corresponding TF can protect DNA from methylation or induce DNA demethylation. Moreover, using the same genetic backbone reduces TF-independent background interference, while using unique molecular barcoding removes bisulfite PCR amplification biases.

### Validation of Hi-TransMet to study the effect of major pioneer transcription factors on DNA methylation at their binding sites

With the aim of identifying factors that can lead to protection from *de novo* methylation or to DNA demethylation, we focused on PFs, as their ability to access nucleosomal DNA in compact chromatin makes them ideal candidates for binding methylated DNA and affecting the methylation status. Twenty-seven PFs were selected based on previously published reports of pioneering activity. Moreover, selected PFs, with the exception of ERα used as a negative control, are expressed in mESCs or in neuronal progenitors (NPs) derived by *in vitro* differentiation of mESCs (**supplementary table 1**)^17^. Consensus wild-type (WT) binding motifs were extracted from public databases^18–20^ or from studies that used ChIP-Seq data for *de novo* motif identification^21, 22^. One binding motif was selected for every TF with a few exceptions: GATA3, 4 and 6 share the same binding motif. SOX2 motif was introduced either alone or in combination with the OCT4 motif as the two factors were shown to often colocalize both in ESCs^23–25^ and during differentiation^26^. Two reported CTCF motifs (CTCF.1 and CTCF.2) corresponding to two different directionalities were included, as the direction of CTCF motif was reported to affect its looping direction^27^ (**supplementary tables 1** and **2**). For each WT motif, we designed a scrambled (Sc) control motif (**supplementary table 2**) with a significantly weaker binding score and in which all CpG positions present in their WT counterparts are maintained. Additionally, we assigned six base pair barcodes on either side of each motif (**figure 1a**). One strand of these barcodes does not contain cytosines and is therefore not affected by bisulfite treatment, thus facilitating motif recognition following bisulfite conversion. We designed the “barcode-motif-barcode” combination by avoiding any resemblance to known TF motifs other than the ones intended (**supplementary table 2**).

**Figure 1.**
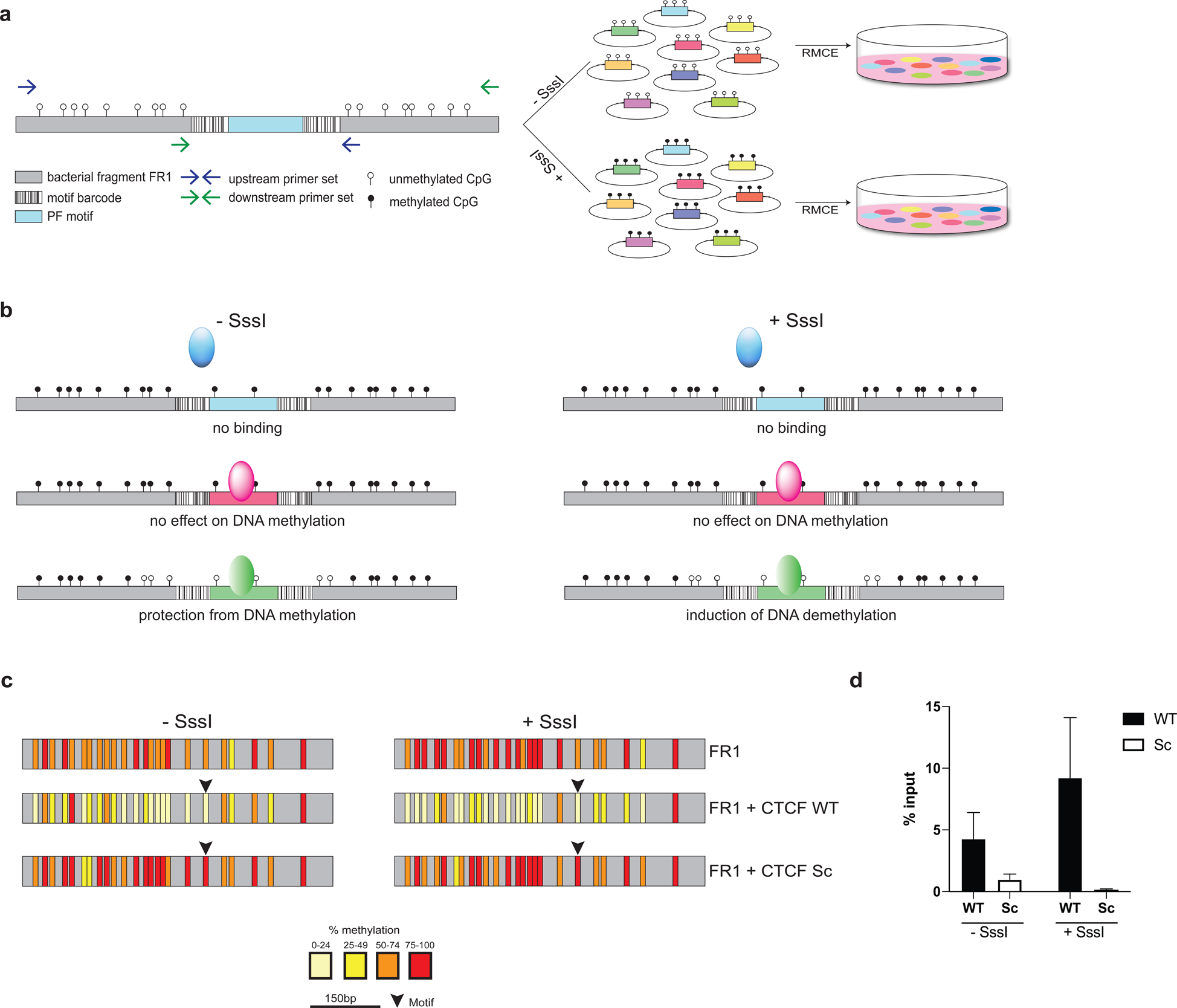
Validation of the experimental approach (Hi-TransMet) to test for TF effect on DNA methylation. (**a**) Schematic representation of Hi-TransMet. PF motifs flanked by unique barcode sequences were individually cloned at the center of an intermediate-CpG content bacterial DNA fragment (FR1) within an RMCE donor plasmid. Motif-containing plasmid libraries were transfected into mESCs following *in vitro* methylation via M.SssI enzyme (+SssI) or without further treatment (–SssI). Insertion of the –SssI library results in *de novo* methylation. Methylation levels in cells that underwent successful recombination were analyzed using universal bisulfite PCR primers designed around the inserted motifs (blue and green arrows). (**b**) Results allow to classify the screened PFs essentially in three classes: I) PFs that cannot bind methylated DNA or induce changes in DNA methylation; II) PFs that are able to bind unmethylated DNA and protect it from methylation; III) PFs that are able to bind methylated DNA and induce DNA demethylation. (**c**) Validation of the experimental approach by Bisulfite Sanger Sequencing in cell lines containing FR1 with CTCF-motifs only. Upon insertion, in the –SssI condition, FR1 undergoes *de novo* methylation. Lower methylation is observed in the presence of the CTCF WT but not Sc motif. In the +SssI condition, high levels of DNA methylation were retained by FR1 in the absence of the motif and in the presence of the CTCF Sc motif. In the presence of the CTCF WT motif, a reduction of DNA methylation levels is observed. Vertical bars correspond to CpG positions, and the color code corresponds to the percentage of methylation calculated for each CpG with a minimum coverage of 10 bisulfite reads. (**d**) CTCF binding in the FR1 was verified by ChIP.

Double-stranded DNA oligomers each representing a unique motif-barcode combination were individually cloned into the FR1 within the RMCE donor plasmid (**supplementary figure 1b**). Plasmids, each containing FR1, one motif and one barcode, were mixed equimolarly to generate the targeting library that was further divided into 2 libraries: one was *in vitro* methylated (+ SssI), while the second did not undergo any further treatment (-SssI). Libraries were separately transfected into the transgenic mESCs, together with a plasmid expressing the CRE recombinase (**figure 1a**). Genomic DNA was then extracted from successfully recombinant cells and treated with sodium bisulfite, followed by amplification of the RMCE site using an approach that amplifies the site regardless of the identity of the inserted motif and labels each molecule with a unique molecular identifier (UMI)^28^. This allows quantifying the methylation of the original unamplified sequences exclusively (**supplementary figure 2a** and **2b**). Reduction of methylation at CpGs surrounding WT binding motifs in comparison to those around the corresponding Sc motifs allows to identify PFs protecting DNA from methylation (–SssI condition) or inducing DNA demethylation (+SssI condition) (**figure 1b**).

**Figure 2.**
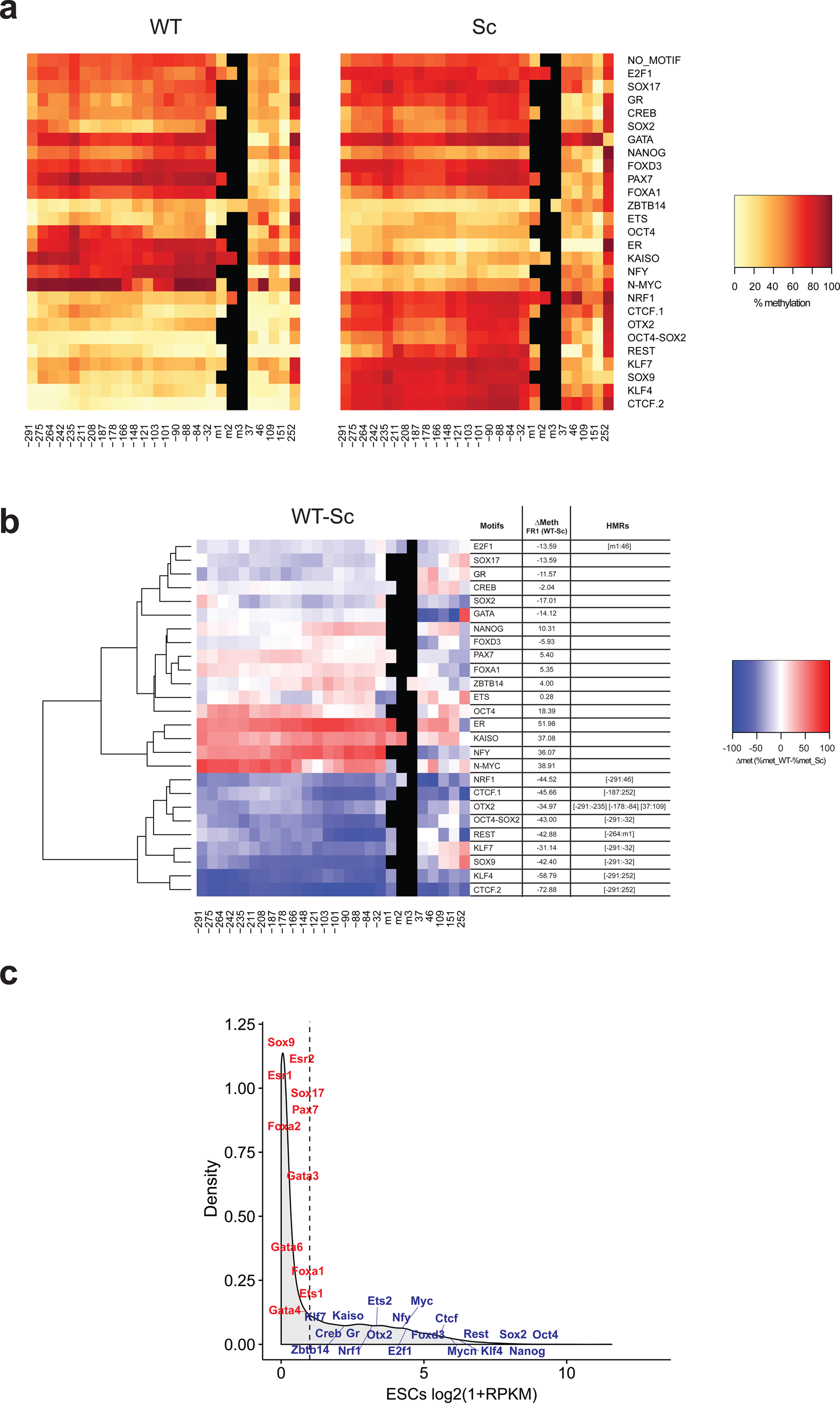
Identification of protective pioneer factors (PPFs) (**a**) Heatmaps indicating methylation percentages of individual CpGs in the FR1 containing WT (left panel) and Sc (right panel) motifs in the –SssI condition. Each line represents a fragment containing the indicated motif. Each square within the line corresponds to one CpG. The methylation percentage of individual CpGs is represented in a color-code. CpGs’ distance from the 5’ end of the motifs is indicated below the heatmaps. CpGs in the motif, when present, are indicated as m1, m2 and m3. (**b**) Differential methylation between WT and Sc motifs in the FR1/–SssI condition. Differential methylation was calculated for each CpG as Δmet = % met_WT – % met_Sc and represented in a color-code. Results were hierarchically clustered using the complete linkage method with Euclidian distance. CpGs’ distance from the 5’ end of the motifs is indicated below the heatmaps. The coordinates of statistically significant hypomethylated regions (HMRs) in WT condition are indicated on the side. (**c**) RNA-Seq density plot of expression levels of the tested PFs in mESCs. A cut-off of Log2(1+RPKM)<1 (dashed line) was used to separate PF expression levels into low (red) and high (blue).

To verify the functionality of Hi-TransMet and its ability to correctly identify factors whose putative binding could lead to lower methylation, we checked the methylation levels of the FR1 fragment including CTCF.1 WT and Sc motifs, as a similar experimental setting was used to study CTCF-mediated DNA demethylation^14^. For this experiment, we used transgenic ESC lines that include FR1 with individual CTCF motifs. The results show that CTCF binding can both protect unmethylated sites from acquisition of *de novo* methylation (-SssI condition) and induce demethylation at methylated sites (+SssI condition). CTCF binding at WT motifs and its absence at Sc motifs were verified by chromatin immunoprecipitation (ChIP) (**figure 1d**). These results validate the use of Hi-TransMet to identify factors that protect DNA from *de novo* methylation or induce DNA demethylation.

### Identification of pioneer transcription factors that can protect DNA against *de novo* methylation at their binding sites

To identify factors that can protect DNA from *de novo* methylation, we analysed reads generated by Hi-TransMet performed under -SssI condition in ESCs. After sorting reads according to their motif-barcode combination, methylation percentages was extracted for all CpGs 300 bp upstream and 250 bp downstream of the inserted motifs **(supplementary figure 2b**). Methylation levels around WT and Sc motifs were first independently analysed (**figure 2a**). As previously reported^12^, the fragments become methylated upon insertion, although not completely, with an average CG methylation level of 52.4% around Sc motifs (**figure 2a** and **supplementary figure 3**). Moreover, while lower methylation levels are observed mostly around WT motifs, there are also changes related to Sc motifs and therefore unrelated to the binding of tested PFs. To correct for this, we subtract, for every motif, the methylation level of each CpG in the locus with the Sc motif from that of the same CpG in the locus with the WT motif (Δmet = % met_WT – % met_Sc, **figure 2b**). Only significantly lower methylation in WT than in Sc conditions will be considered as directly related to PF binding. We then classified different effects of TF motifs on DNA methylation using unsupervised hierarchical clustering, followed by the identification of statistically significant hypomethylated regions in WT conditions (HMRs). HMRs are defined as regions of more than 50bp and containing a minimum of 3 consecutive CpGs, each having a Δmet of 10% or higher.

**Figure 3.**
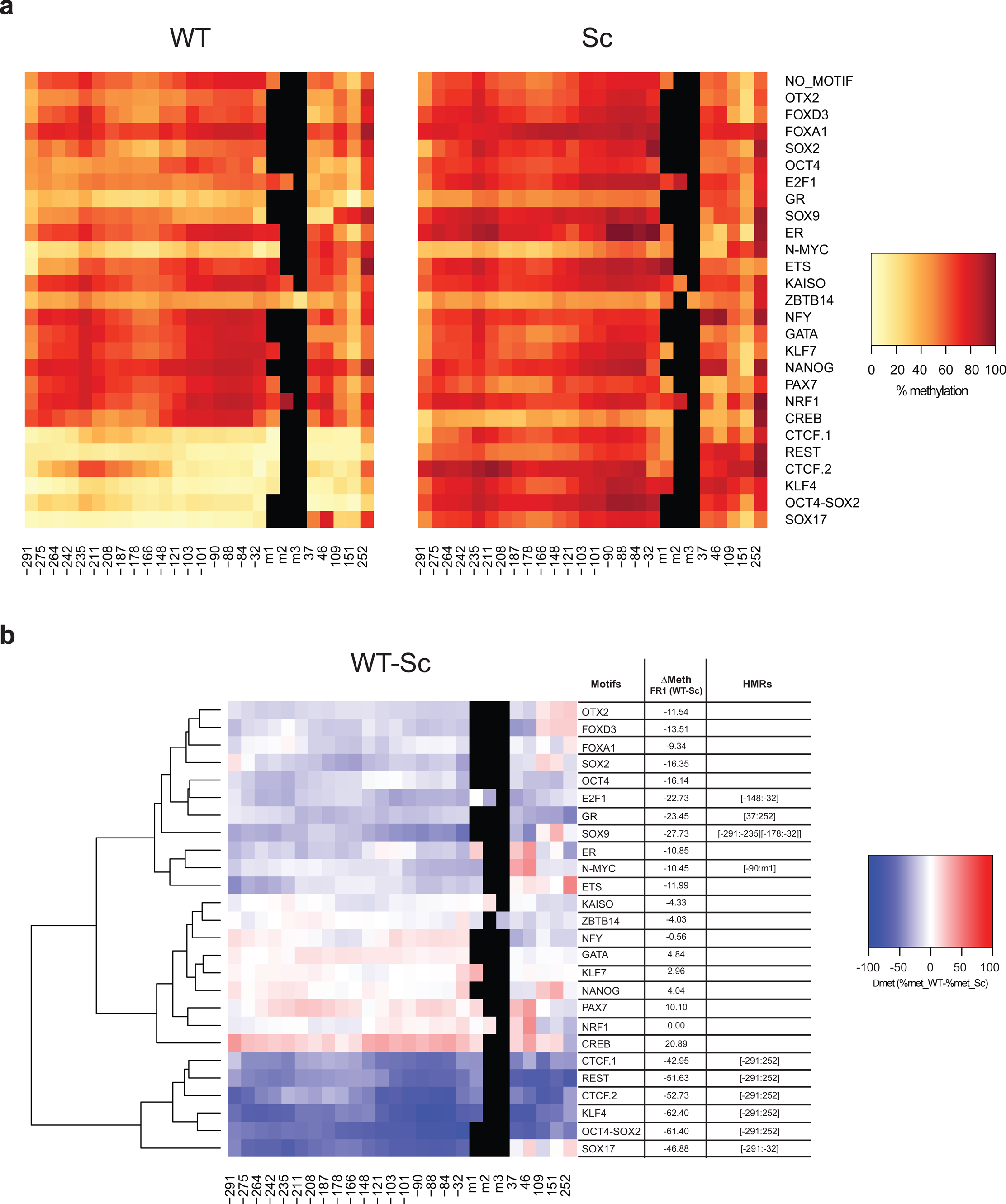
Identification of super pioneer transcription factors (SPFs) (**a**) Heatmaps indicating methylation percentages of individual CpGs in the FR1 containing WT (left panel) and Sc (right panel) motifs in the +SssI condition. Each line represents a fragment containing the indicated motif. Each square within the line corresponds to one CpG. The methylation percentage of individual CpGs is represented in a color-code. CpGs’ distance from the 5’ end of the motifs is indicated below the heatmaps. CpGs in the motif, when present, are indicated as m1, m2 and m3. (**b**) Differential methylation between WT and Sc motifs in the FR1/+SssI condition. Differential methylation was calculated for each CpG as Δmet = % met_WT – % met_Sc and represented in a color-code. Results were hierarchically clustered using the complete linkage method with Euclidian distance. CpGs’ distance from the 5’ end of the motifs is indicated below the heatmaps. The coordinates of statistically significant hypomethylated regions (HMRs) in WT condition are indicated on the side.

Results show that PFs differ in their ability to protect DNA from methylation. Globally, it seems that, upon binding, only few of the selected PFs can protect from *de novo* DNA methylation. We called these “protective pioneer factors (PPFs)”. In addition to the previously reported^14, 30^ CTCF (CTCF.1 and CTCF.2) and NRF1, our results indicate that also KLF4, KLF7, OCT4-SOX2, SOX9, REST, OTX2, and E2F1 protect against methylation (**figure 2b**). Moreover, SOX2 alone, but not OCT4, seems to be able to protect against methylation, although this ability is increased in the presence of a combined OCT4-SOX2 motif. It is important to note that all identified PPFs, with the exception of SOX9, are highly expressed in ESCs (**figure 2c**).

### Identification of super pioneer transcription factors that can induce DNA demethylation at their binding sites

To identify PFs that can cause DNA demethylation upon binding to methylated DNA in ESCs, we analysed methylation levels in the +SssI condition. The inserted fragment maintained high levels of methylation in ESCs, with an average CG methylation level of 79.1% in fragments with Sc motifs (**figure 3a** and **supplementary figure 3a**). Under these conditions, we observed extensive DNA demethylation around the binding sites of several factors: CTCF (CTCF.1 and CTCF.2), REST, KLF4, OCT4-SOX2, SOX9, SOX17, E2F1, N-MYC, and GR. Moreover, there is again a considerable reduction of DNA methylation around the SOX2 motif, while this is less apparent around OCT4 motif (**figure 3b**). We called the corresponding factors “super pioneer transcription factors (SPFs)” as, in addition to their known ability to induce chromatin remodelling, they are also able to induce DNA demethylation (**figure 2c**). It is interesting to note that CTCF, REST, SOX2, SOX9, E2F1 and KLF4 TFs both protect from *de novo* methylation and induce DNA demethylation. On the other hand, NRF1 and OTX2, can only protect DNA from methylation but have no effect on methylated DNA. This is in concordance with a previously published study defining NRF1 as methylation sensitive^30^. Similar to PPFs, most SPFs, with the exception of SOX9 and SOX17, are highly expressed in ESCs.

Interestingly, clustering of the results revealed that reduction in DNA methylation at some PPF and SPF binding sites extends far beyond the TF binding site. This could be due to the sequence context of our reporter DNA fragment, which lacks motifs for other TFs, but might also suggest more active mechanisms, rather than steric hindrance, used by PFs to maintain low levels of DNA methylation and render a large region available for the binding of settler TFs.

### PPFs and SPFs are cell-type specific

PPF- and SPF-mediated effects on DNA methylation are expected to be dynamic during differentiation as a function of cell-type-specific TF expression. Therefore, we differentiated the transgenic ESCs into neuronal progenitors (NPs)^17^, in order to both confirm our results in ESCs and identify new NP-specific PPFs and SPFs.

Comparison of gene expression profiles derived by RNA-Seq data in ESCs and NPs highlighted the differences in expression of the tested PFs between the two cell types (**figures 4a** and **4b**). Hi-TransMet was then performed in NPs and methylation levels around PF motifs in ESCs and NPs were compared. First, differential expression of each tested PF in ESCs and NPs was plotted against the difference in Δmet between ESCs and NPs (ΔΔmet = Δmet ESCs - Δmet NPs) of the FR1 containing the corresponding PF motif. This showed an overall anticorrelation in both -SssI and +SssI conditions (**figures 4c** and **4d**), indicating that most methylation changes are indeed driven by the direct activity of the corresponding PFs.

**Figure 4.**
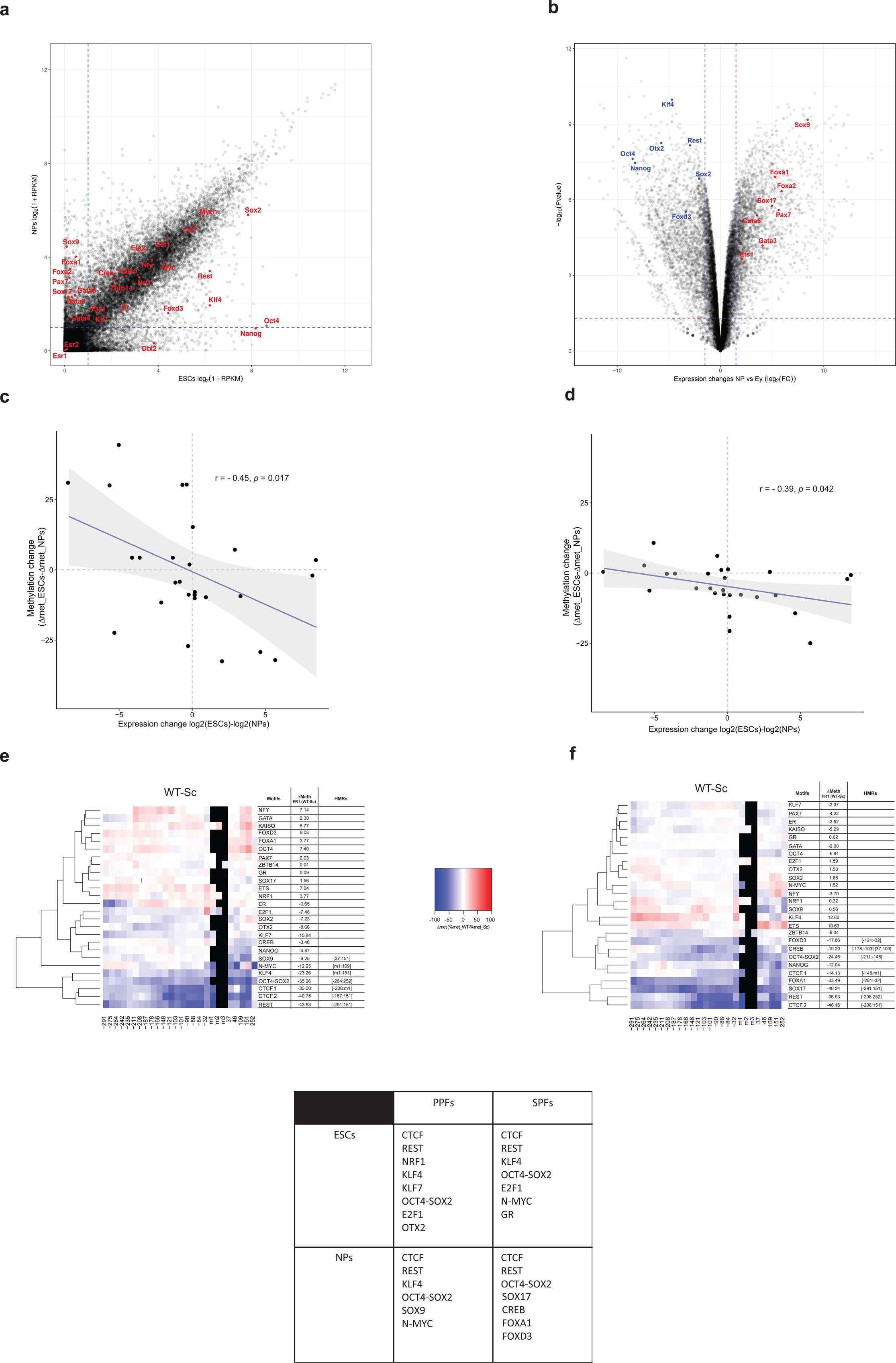
PPFs and SPFs are cell-type specific. (**a**) Scatter plot showing differential gene expression between ESCs (x axis) and NPs (y axis) based on RNA-Seq data. Tested PFs are labeled in red. A cut-off of Log2(1+RPKM)<1 (dashed lines) was used to separate PF expression levels into low and high. (**b**) Volcano plot highlighting genes with a significantly different expression levels between ESCs and NPs. Cut-off indicated by the dashed lines. (**c**,**d**) Scatter plots comparing differential expression of each tested PF in ESCs and NPs against the difference in Δmet between ESCs and NPs (ΔΔmet = Δmet_ESCs - Δmet_NPs) of the FR1 containing the corresponding PF motif in -SssI (**c**) and +SssI (**d**) conditions. Each dot represents changes in average Δmet of FR1 fragment with one motif. r= Pearson correlation coefficient; p= p-value. (**e**,**f**) Differential methylation (Δmet) between WT and Sc motifs in the FR1/–SssI (**e**) and in the FR1/+SssI (**f**) conditions in NPs. Differential methylation was calculated for each CpG as Δmet= %met_WT – %met_Sc and represented in a color-code. Results were hierarchically clustered using the complete linkage method with Euclidian distance (see materials and methods). CpGs’ distance from the 5’ end of the motifs is indicated below the heatmaps. The coordinates of statistically significative hypomethylated regions (HMRs) in WT condition are indicated on the side. (**g**) Table summarizing identified PPFs and SPFs in ESCs and NPs.

Average methylation levels highly increased during differentiation, reaching 81.7% (Sc motifs) in the -SssI condition (**supplementary figures 3a** and **4a**) and 85.9% in the +SssI condition (**supplementary figures 3a** and **4b**). In the -SssI condition, statistical analysis identified HMRs around CTCF.1, CTCF. 2, REST, KLF4, OCT4-SOX2, SOX9 and N-MYC binding sites (**figure 4e**). On the other hand, FR1/+SssI data analysis identified CTCF, REST, OCT4-SOX2, SOX17, CREB, FOXA1 and FOXD3 as SPFs (**figures 4f** and **4g**).

### Most SPFs induce TET-dependent DNA demethylation

Next, we sought to determine the mechanisms used by SPFs to demethylate their binding sites. DNA demethylation could occur in a replication-dependent fashion through the inhibition of the methylation maintenance machinery, notably the DNA methyltransferase DNMT1. Another possibility is the SPF-dependent induction of replication-independent active demethylation processes.

Ten-Eleven Translocation (TET)-dependent oxidation of 5-methylcytosine (5-mC) into 5-hydroxymethylcytosine (5-hmC) is currently considered as an essential step for active DNA demethylation. Several groups published interactions between PFs and TET enzymes^31–37^ (**supplementary table 1**), consistent with studies reporting correlation between low levels of 5mC and high levels of 5hmC and TET proteins at TF-binding sites^31, 38^.

To address the functional involvement of TET proteins in SPF-dependent DNA demethylation, we performed Hi-TransMet on FR1/+SssI in mESCs lacking all TET proteins: TET1/2/3 triple knockout (TKO)^39^, to study the capacity of the identified SPFs to induce DNA demethylation in this context. In the absence of TET proteins, average methylation levels of FR1 are significantly higher than in ESCs expressing TETs, both in CG context (88.6% at Sc motifs, **figure 5a** and **supplementary figure 3a**) and non-CG context (6.3%, **supplementary figure 3b**), suggesting that TET proteins are responsible for most binding-specific and unspecific demethylation events observed in the previous experiments. Results show that most SPF-dependent DNA demethylation activity is weak or absent in TET TKO cells, indicating that most SPFs induce active DNA demethylation (**figure 5b**).

**Figure 5.**
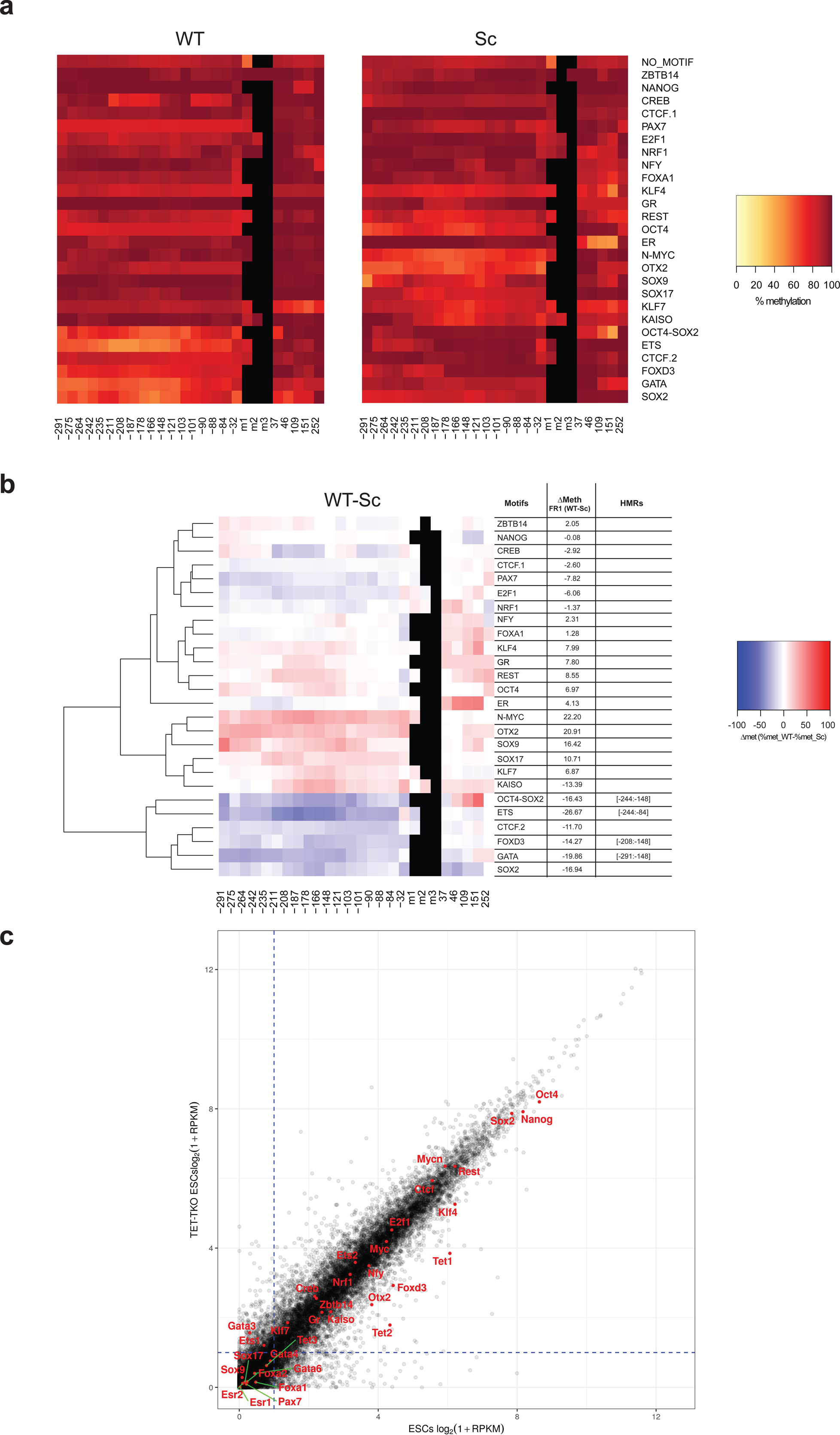
Most SPFs induce TET-dependent active DNA demethylation. (**a**) Heatmaps indicating methylation percentages of individual CpGs in the FR1 containing WT (left panel) and Sc (right panel) motifs in the +SssI condition in TET TKO ESCs, as in figures 2a and 3a. (**b**) Differential methylation between WT and Sc motifs in the FR1/+SssI condition in TET TKO ESCs, as in previous figures. (**c**) Scatter plot showing differential gene expression between WT (x axis) and TET TKO (y axis) ESCs based on RNA-Seq data. Tested PFs as well as TET genes are labeled in red. A cut-off of Log2(1+RPKM)>1 (dashed lines) was used to separate PF expression levels into low and high.

Interestingly, demethylation occurs in the absence of TETs at the OCT4-SOX2 binding site. This is also observed at the SOX2 motif alone, although no statistically significant HMRs were identified. Other PF motifs that have lower methylation under these conditions are FOXD3, GATA and ETS. GATA factors and ETS have very low expression in ESCs although they are slightly upregulated in TET TKO cells (**figure 5c**). It is therefore unlikely that the effect we see around their corresponding motifs is directly driven by these factors. On the other hand, FOXD3 is both highly expressed in TET TKO cells (**figure 5c**) and shows moderate SPF activity in NPs. SOX2, and FOXD3 might therefore lead to passive DNA demethylation. As TET TKO ESCs cannot differentiated into NPs, NP-specific SPFs as well were included in the following experiments aimed at testing SPF-dependent passive demethylation.

### SOX2 inhibits DNMT1 activity

As maintenance of DNA methylation is catalysed by DNMT1, we set up an *in vitro* methylation assay to assess the effect of PFs on DNMT1 activity^40^. A double stranded hemi-methylated DNA probe containing the PF motif of interest and a single CpG (either within or in the immediate vicinity of the motif) was incubated with DNMT1 protein and radioactively-labelled S-Adenosyl-L-methionine (SAM[3H]) as a methyl donor, in the presence or absence of the corresponding PF. Integration of the radioactively-labelled methyl group in the unmethylated strand was measured as a readout of DNMT1 activity and for each PF, the signal in the presence of the WT motif was normalized to the signal in the presence of the Sc motif.

Surprisingly, results showed that only SOX2, and to a lesser extent OTX2 and ETS, among all tested SPFs and non-SPFs, significantly reduce DNMT1 activity (**figure 6a**). Moreover, the presence of SOX2 alone, but not OCT4, is sufficient to significantly reduce DNMT1 activity on the combined OCT4-SOX2 probe, further confirming that SOX2 inhibits DNMT1 activity on hemi-methylated DNA (**figure 6b**). Unlike SOX2, FOXD3 does not affect DNMT1 activity. This suggests that FOXD3 might induce a TET-independent active demethylation. Finally, NP-specific SPFs SOX17, CREB, and FOXA1 do not affect DNMT1 activity, suggesting that they depend on TETs to induce demethylation.

**Figure 6.**
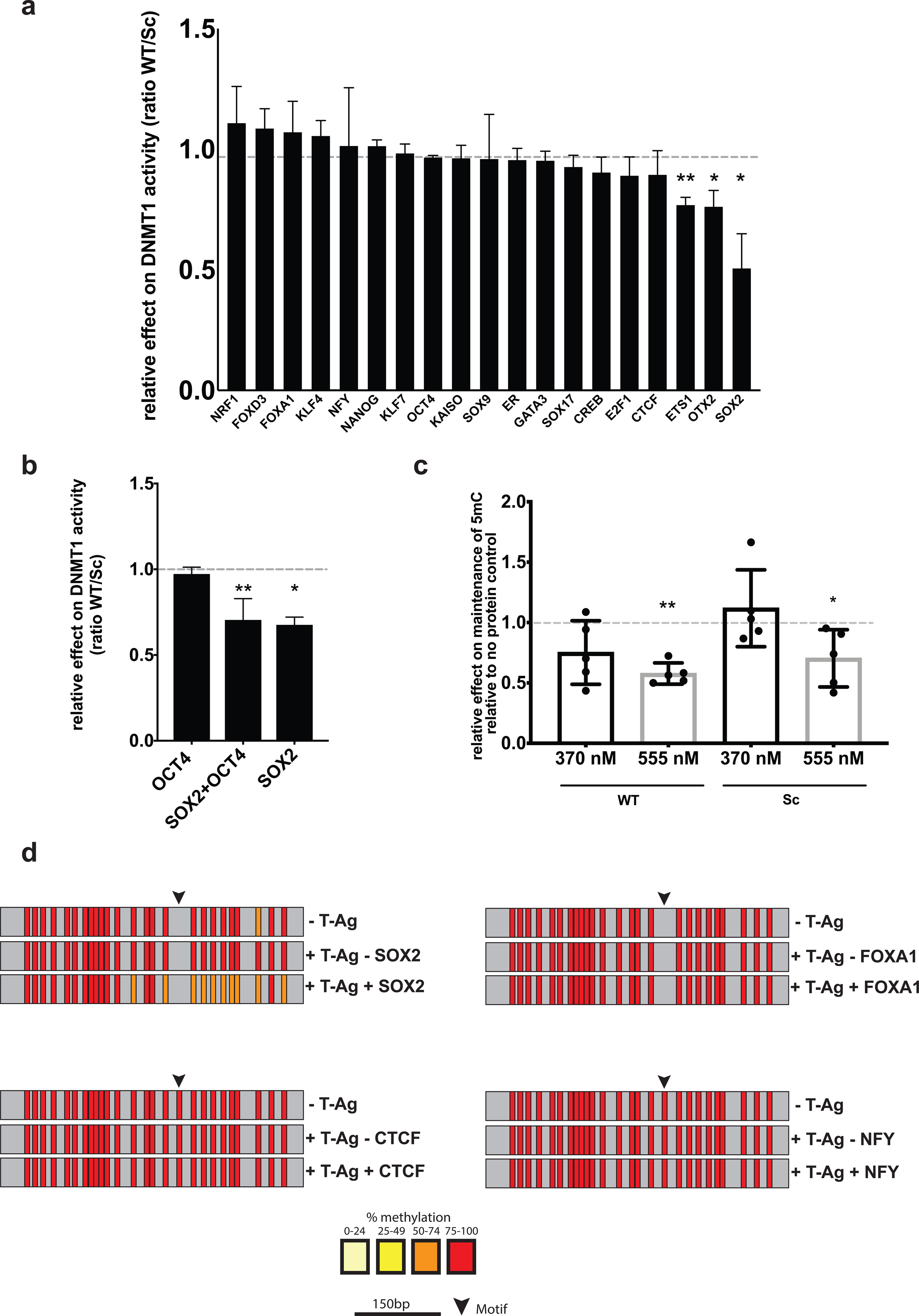
SOX2 inhibits DNMT1-dependent maintenance of DNA methylation during replication. (**a**) *In vitro* methylation assay to measure DNMT1-mediated DNA methylation using hemi-methylated probes containing WT or Sc PF motifs in the presence or absence of the corresponding PF. Relative DNMT1 activity is represented as scintillation counts, corrected for the weight of isolated DNA probes and compared to the Sc probe. Results are shown as mean + SEM of three biological triplicates. SOX2, ETS and OTX2 significantly reduce DNMT1 activity (ETS1 p= 0.0018, OTX2 p=0.0238, SOX2 p= 0.026, two-tailed unpaired t-test). **b**) *In vitro* methylation assay using the OCT4-SOX2 motifs in the presence of SOX2 alone, SOX2+OCT4 and OCT4 alone. Results are shown as mean + SEM of three biological triplicates (SOX2+OCT4 p= 0.0035, SOX2 p=0.0185, two-tailed unpaired t-test). **c**) *In vitro* replication assay to assess the effect of TF binding on the maintenance of DNA methylation during replication. Methylation levels following replication are measured based on the integration of radioactively labeled methyl group during replication. Results are presented as mean ± SD of five biological replicates and analyzed as radioactive signal in the presence of SOX2 relative to the signal in absence of SOX2. P values: SOX2_WT 370nM p=0.1026, 555nM p=0.0004; SOX2_Sc 370nM p=0.4521, 555nM=0.0491, two-tailed unpaired t-test. **d**) Bisulfite Sanger sequencing analysis of a bacterial DNA fragment before or after replication (±T-Ag) and in the absence or presence of the PF (± PF). Vertical bars correspond to CpG positions, and the color code corresponds to the percentage of methylation calculated for each CpG with a minimum coverage of 10 bisulfite reads.

### SOX2 inhibits DNA methylation maintenance during replication

To address if SOX2-dependent inhibition of DNMT1 takes place during DNA replication, we setup an *in vitro* replication assay^41, 42^ that assesses the effect of TFs on the maintenance of DNA methylation. Briefly, a bacterial DNA fragment containing the tested motif is cloned into an SV40 replication vector^43^ to generate the replication substrate. Incubation of the substrate with T-Antigen (T-Ag) and cellular extracts leads to its replication. Addition of biotinylated dUTP to the reaction results in biotinylation of nascent DNA. Thus, Immunoprecipitation with streptavidin beads enriches for newly synthesized biotinylated DNA. Complete replication is verified by digestion with DpnI, an enzyme that cuts specifically at GATC sites when the Adenosine is methylated. As m6A is not maintained during replication, replicated templates are protected from digestion (**supplementary figure 5b**). Finally, the substrate could be *in vitro* methylated by treatment with SssI prior to the replication reaction, and the maintenance of methylation can be assessed by adding SAM[3H] to the replication reaction and counting the integration events of the radioactive methyl group or by bisulfite sequencing of the replicated product.

We performed the replication reaction in the presence or absence of the PF of interest. PF binding to the plasmid was verified by EMSA (**supplementary figure 5c**), and SAM[3H] incorporation was measured by scintillation counting and normalized to the signal in absence of PFs. Results show that the substrate replicates efficiently (**supplementary figure 5d**) and replicated templates maintain DNA methylation in the absence of PFs (**supplementary figure 5d** and **figure 6d**). Unmethylated templates, and templates incubated in the absence of T-Ag (which do not replicate) do not incorporate SAM[3H], confirming that SAM[3H] incorporation reflects maintenance methylation. Interestingly, in the presence of SOX2 protein, we observed a significant reduction in methyl group incorporation around the WT motif, clearly suggesting that SOX2 interferes with the maintenance of DNA methylation during replication. Moreover, analysis of the methylation status around the motifs by bisulfite Sanger sequencing of replicated DNA in two independent replicates (**figure 6d** and **supplementary figure 5e**) confirmed a reduction in DNA methylation in the presence of SOX2, but not of CTCF, FOXA1 and NFY, further suggesting that SOX2 recruitment leads to passive DNA demethylation by DNMT1 inhibition.

## Discussion

While PF effect on nucleosome compaction is well documented, PF interaction with DNA methylation is still poorly addressed. Here, we established Hi-TransMet, a high-throughput approach to assess the effect of TFs on DNA methylation. While we focused on PF crosstalk with DNA methylation, this method could be used with any DNA-binding factor of interest and the throughput can be easily increased.

It is important to note that all WT motifs used here were previously tested experimentally for their ability to specifically and efficiently recruit their corresponding TFs, either by ChIP experiments, or DNA/protein microarrays and EMSA. Therefore, we made the assumption that the TFs bind to the WT motifs in our setting and therefore that lower DNA methylation levels around WT motifs in comparison to those around Sc motifs are linked to this binding event. This assumption was validated by ChIP assays performed on selected PFs (**supplementary figure 6**). Using Hi-TransMet, we identified PPFs that are able to protect against *de novo* methylation. Our screening both confirms previously reported PPFs (NRF1^30, 44^, CTCF and REST^14^) and identifies new ones, either constitutive (KLF4, SOX2, SOX9) or ESCs-(KLF7, E2F1 and OTX2) and NP-specific (N-MYC) (**figure 4g**). Whether PPF binding shields its surrounding from DNA methyltransferases by steric hindrance or whether PPFs directly interact with DNMT3a/3b/3L leading to their inhibition awaits further studies. We also identified SPFs that, in addition to their known pioneering activities, can induce DNA demethylation at their binding sites. Constitutive SPFs are CTCF, REST, SOX2, and SOX17. ESC-specific SPFs are KLF4, E2F1, GR, N-MYC and SOX9 while NP-specific SPFs are FOXA1, FOXD3, and CREB (**figure 4g**).

Considering the previously established PF ability to access their binding sites in a closed chromatin context, the identification of SPFs introduces a further level of classification and suggests a clear hierarchy among TFs in the fine regulation of gene expression. We propose a model where SPFs are the first to engage methylated binding sites. This is followed by DNA demethylation allowing the recruitment of “normal” PFs and further chromatin opening, which provides access to settler TFs (**figure 7**).

**Figure 7.**
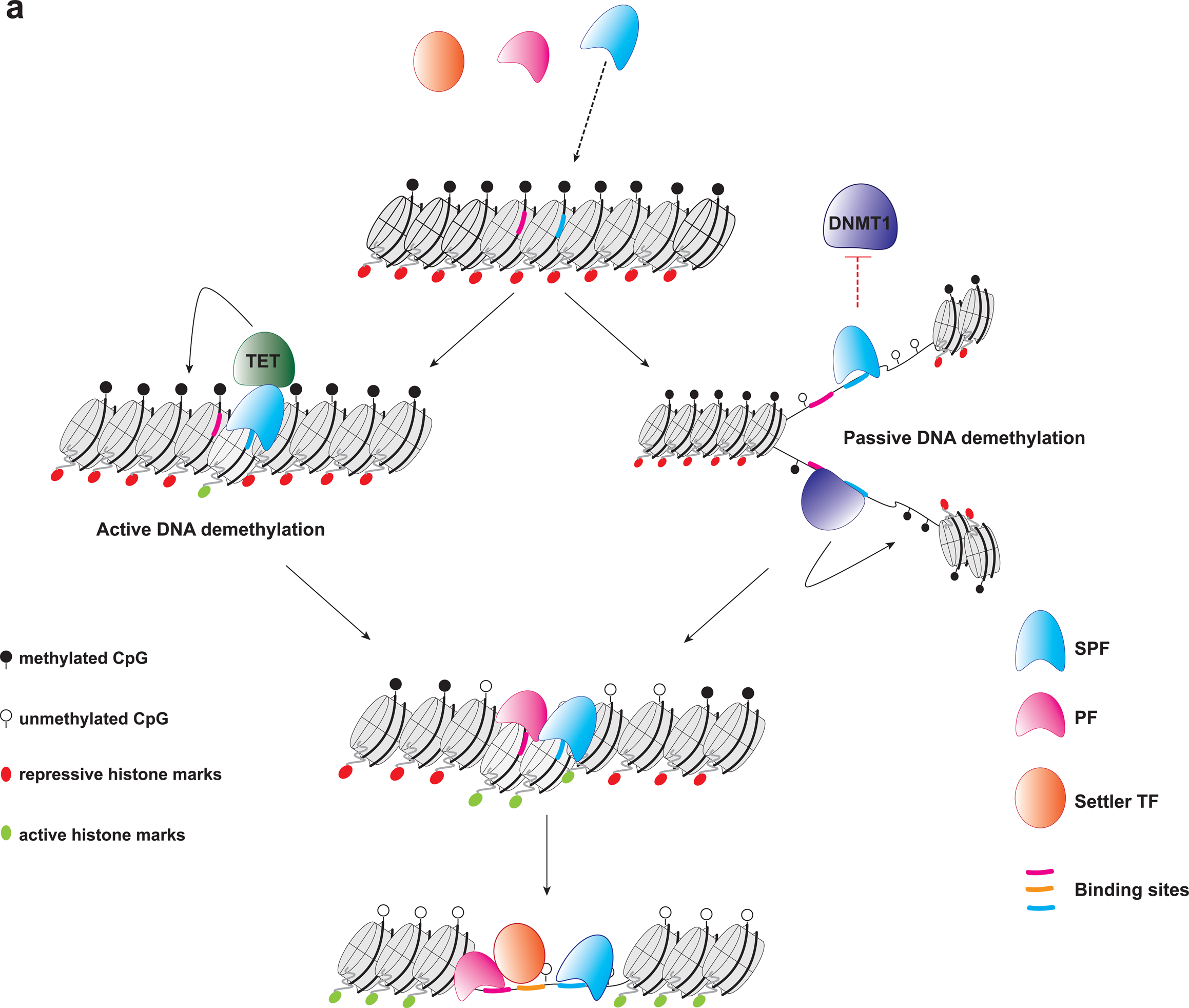
Hierarchy of Transcription Factor binding. (**a**) super pioneer transcription factors (SPFs) engage their target sequences in closed chromatin and in the presence of DNA methylation. Upon binding, most SPFs drive DNA demethylation through active processes, mainly mediated by the TET enzymes, whereas SOX2 leads to passive DNA demethylation. Loss of DNA methylation allows the binding of methylation sensitive PFs. Nucleosome remodelling and deposition of histone modifications associated with open chromatin regions, mediated by both SPFs and PFs, create a favorable environment for the binding of “settler” TFs.

In accordance with our results, CTCF and REST were both predicted to induce DNA demethylation at their binding sites^14^. Similarly, several FOX factors were linked with DNA demethylation and TET1^45–48^. Moreover, overexpression of FOXA2, a paralog of FOXA1, in fibroblasts correlates with chromatin opening and loss of methylation at its target sites. The pluripotency factor KLF4 was also recently shown to mediate active DNA demethylation at closed chromatin regions by interacting with TET2 during reprogramming^31^. Conversely, KLF7 was shown to interact with DNMT3a in TF protein array studies^49, 50^.

PPF and SPF activity seems to depend not only on their expression levels, but also on their interactors, PTMs, and roles in the different cell lines. For example, retinoic acid (RA)-mediated NP differentiation drives CREB phosphorylation, a necessary modification for its DNA binding ability, which could explain its NP-specific SPF activity^51, 52^, despite its expression in both ESCs and NPs (**supplementary figure 6**). While several studies have proposed CREB to be sensitive to DNA methylation^53, 54^, the lack of CREB phosphorylation in the assays used in those studies may explain this discrepancy. Also, although expressed at similar levels, NRF1 and KLF7 lose their protecting ability in NPs. This might indicate a lower efficiency in stably protecting against methylation in conditions where higher levels of DNA methylation are the default status, as it the case in NPs. N-MYC, on the other hand, induces DNA demethylation in ESCs, but only protects against methylation in NPs, suggesting that it is unable to demethylate highly methylated regions. Previous reports on N-MYC are contradictory: on one hand, N-MYC was shown to be strongly linked to regions harbouring H3K4me3 histone marks, therefore most likely having low DNA methylation levels^55^, and loss of N-MYC was associated with heterochromatinization in neuronal stem cells^56^. On the other hand, N-MYC was reported to bind to hypermethylated regions in neuroblastoma cell lines, although binding sites in this study were depleted of the E-box CACGTG that was used in our study^57^. While GR was previously shown to bind methylated cytosines in non-CG context, its effect on DNA methylation was not assessed^58^. Finally, SOX9 and SOX17 have low expression levels in ESCs, but seem to behave as PPFs and SPFs in these cells: while low expression levels may be sufficient for this activity, it cannot be excluded that the motifs chosen might also be recognized and bound by other SOX family members.

OCT4 and SOX2 are known to widely colocalize in the genome of ESCs^26, 59, 60^, and were reported to be involved in maintaining a hypomethylated state at the maternal *Igf2/H19* ICR, possibly through DNA demethylation^61, 62^, or by protection from *de novo* methylation^63^, although a mechanism of action was not formally proposed. In our study SOX2 shows a tendency, to act as a PPF, in both cell types, and as a SPF, in ESCs. It can be hypothesized that, while SOX2 is the factor involved in mediating protection from methylation and demethylation, OCT4 might stabilize SOX2 binding, thus amplifying the effect on DNA methylation. Accordingly, it was recently shown that OCT4 might not be essential for the generation of iPSCs from fibroblasts, pointing towards a higher ranking for SOX2 in the hierarchy of OKSM activity^64^. In NPs, OCT4 is silenced and replaced by the related POU family member BRN2 (POU3F2) in its interaction with SOX2^24^. Interestingly, the BRN2 binding motif is highly similar to that of OCT4, so it is plausible to hypothesize that the SOX2-BRN2 interaction in NPs has a similar effect on DNA methylation.

Importantly, we show that SOX2 mediates replication-dependent passive demethylation. Although the need for replication was reported for several TFs to induce DNA demethylation^111^, this is, to our knowledge, the first evidence of direct TF interference with the activity of DNMT1 during replication, in mammals. However, the exact mechanisms by which SOX2 mediates such an effect are yet to be elucidated. Based on the current knowledge, two possible mechanisms of SOX2-mediated passive DNA demethylation can be hypothesized:1) SOX2 binding at the replication fork inhibits DNMT1 activity by steric hindrance. In this model, SOX2 binding would precede DNMT1 recruitment, as it was demonstrated that there is a delay in the recruitment of DNMT1 to the replication fork^65, 66^. 2) SOX2 directly interacts with and inhibits components of the maintenance machinery. Indeed, a weak interaction between UHRF1 and SOX2 has been reported^67^. Finally, it would be interesting to assess the extent of this phenomenon and whether it is shared by other TFs. If it is the case, this could constitute another piece of the puzzle explaining the maintenance, or the lack thereof, of epigenetic modifications during replication.

## Online Methods

### Cell culture

TC-1(WT) ES cells^12^ were cultivated on feeder cells or on dishes coated with 0.2% porcine skin gelatin (Sigma, cat. No. G1890) in high glucose-DMEM medium (Gibco^TM^, cat. No. 31966021) supplemented with 1% NAA (Gibco^TM^, 11140035), 1:1000 homemade LIF and 1.42nM beta mercaptoethanol. Differentiation into neuronal progenitors was performed as previously described^17^. Briefly, ESCs were grown for 4 days in CA medium (DMEM, 10% FBS, 1% NAA,1.42nM beta-mercaptoethanol) in non-adherent plates (Greiner, Bio-one 94/16 with vents, 633102), then supplemented with 5μM Retinoic Acid (Sigma, R-2625) for another 4 days. Medium was changed every two days.

### Insertion of RMCE cassette in TET TKO cell lines

Insertion of the RMCE Hy-TK cassette into the TET TKO mouse ESCs was performed as previously described^12^. Briefly, 4×10^6^ cells were transfected with 100µg of the pZRMCE plasmid linearized with SapI (NEB, R0569S) using the Nucleofector^TM^ 2b device and the Mouse ES cells Nucleofector^TM^ kit (Lonza, VAPH-1001). The plasmid includes a 2.4 kb and 3.1 kb homologous arms to the positions - 1300 upstream and +2332 downstream of the *Hbb-γ* ATG start, respectively. These arms flank two inverted *LoxP* sites which, in turn, flank the selection cassette. Upon transfection, positive selection of clones was done using 25μg/mL Hygromycin B Gold (InvivoGen, ant-hg-1) for 12 days. Surviving colonies were picked and screened for successful insertion by PCR (primer sequences in **supplementary table 3**).

### Recombinase Mediated Cassette Exchange (RMCE)

The bacterial fragment FR1 (**supplementary figure 1b**) was synthesized by Invitrogen GeneArt Gene Synthesis and inserted into the RMCE donor plasmid by directional cloning using the restriction enzymes BamHI (NEB, R3136S) and HindIII (NEB, R3104L). Single stranded oligomers containing the motifs were synthesized by ThermoFisher Scientific. For each motif, forward and reverse oligomers were annealed and cloned into the FR1 fragment by directional cloning using the restriction enzymes SphI (NEB, R3133L) and NheI (NEB, R3131L). To create the plasmid libraries, single-motifs containing plasmids were mixed in equimolar fashion and co-precipitated before RMCE or M.SssI treatment.

RMCE transfection was performed as previously described^13^. Cells containing the Hy-TK RMCE cassette were cultured in ES medium (15% FBS) containing 25μg/ml hygromycin for at least 10 days and split the day before transfection. Medium was changed to 20% FBS ES medium 2 hours before electroporation. Cells were then washed with PBS, detached and counted. 4 million cells were electroporated with 75μg of the targeting plasmid or plasmid libraries and 45μg of plc-CRE plasmid and plated in two P10 dishes with 20% FBS ES medium, as before. Positive selection with 3μM ganciclovir (NEB, CLSYN001) was started two days after transfection. After 12 days, surviving colonies were picked and screened for correct insertion via PCR (primer sequences in **supplementary table 3**).

### Plasmid methylation by M.SssI treatment

When indicated, plasmid libraries were methylated before transfection using the M.SssI CpG methyltransferase (NEB, M0226L) as previously described^68^. Briefly, 100μg of plasmids were incubated with 1x NEBuffer 2, 32mM SAM and 22.5μL 20000 units/mLM.SssI for 30 minutes. The reaction was then replenished with the same amounts of SAM and M.SssI in a final volume of 500μL and incubated at 37°C for another hour. Plasmid DNA was purified with phenol-chloroform and precipitated with ethanol. Complete methylation of the samples was verified by digestion with HpaII (NEB, R0171L), a methylation sensitive restriction enzyme, and methylation unsensitive MspI (NEB, R0106L) as a control.

### Bisulfite conversion and PCR

Genomic DNA was extracted using the GenElute Mammalian genomic DNA miniprep kit (Sigma, G1N70-1KT). Bisulfite conversion of 800ng gDNA (for Sanger sequencing) or 3μg (for Hi-TransMet library preparation) was conducted using the EZ DNA Methylation-Gold™ Kit (Zymo Research, D5006). Regions of interest were amplified by PCR using the AmpliTaq Gold^TM^ DNA Polymerase (Applied Biosystems^TM^, N8080241) and ran on a 1% agarose gel. Bisulfite PCR program: 95°C 15 min; 20 touch-down cycles from 61 to 51°C with 30 sec at 95°C, 30 sec annealing T and 1 min at 72°C; 40 cycles of 30 sec at 95°C, 30 sec at 53°C and 1 min at 72°C; final extension at 72°C 15 min.

For Sanger sequencing, PCR products were extracted from 1% Agarose gels using the GenElute Gel Extraction kit (Sigma, NA1111-1KT) and cloned into the pCR^TM^4-TOPO plasmid of the TOPO® TA Cloning® Kit for Sequencing (Invitrogen, K45750), transformed into TOP10 bacteria, and plated on Agar dishes with 100μg/mL Ampicillin. Individual bacterial colonies were picked, followed by amplification and DNA extraction using the GenElute^TM^ HP Plasmid Miniprep Kit (Sigma, NA0150-1KT). Finally, the products were sequenced using the M13r primer. Results were analysed using the BISMA or BiQ Analyzer online tools (http://services.ibc.uni-stuttgart.de/BDPC/BISMA/ and https://biq-analyzer.bioinf.mpi-inf.mpg.de/)^69,70^.

### Hi-TransMet Library Preparation and Sequencing

The UMI-based library protocol consists of 3 steps: annealing, non-barcoded amplification and adapters addition (**supplementary figure 2a**). For each library, 3μg of bisulfite converted DNA were used as starting material. Annealing program: 95°C 15 min; gradual temperature decrease from 61°C to 51°C, −0.5°C/min; final extension at 72°C 7 min. Annealing was performed using the AmpliTaq Gold^TM^ DNA Polymerase (Applied Biosystems^TM^, N8080241), reaction set up according to the manufacturer’s protocol. Following a purification step to remove unused primers, annealed DNA was subjected to a short amplification with a universal forward primer and a specific reverse bisulfite primer: 95°C 10 min; 3 cycles of 95°C 15 sec, 50°C 30 sec, 72°C 1 min; final extension 72°C 5 min. Amplified DNA was purified of the reaction mix, then sequencing adapters were added in a final amplification step: 95°C 15 min; 30 cycles of 95°C 15 sec and 60°C2 min. Primer dimers were eliminated in a final purification step. Library barcodes and primers are listed in **supplementary tables 4** and **5**. The PCR steps were done using the Promega Go-TaqG2 Hot Start Green Mastermix (Promega, M7423), set up according to manufacturer’s protocol, and all the purification steps using the Qiagen GenRead Size Selection kit (Qiagen, 180514). Correct library size was verified using the Agilent 2200 Tape Station system (Agilent, G2964AA, 5067-5584 and 5067-5585). Libraries were sequenced using Illumina MiSeq platform generating 300 base pair paired-end reads (PE300).

### Chromatin Immunoprecipitation (ChIP)

ChIP was performed using the Diagenode IP-Star Compact Automated System robot (Diagenode, B03000002) and the Diagenode AutoiDeal ChIP-qPCR kit standard protocol (Diagenode, C01010181) on 4×10^6^ cells. Sonication was performed using the Diagenode Bioruptor Pico (Diagenode, B01060010) and the following conditions: 8 cycles of 30 sec ON and 30 sec OFF for mESCs; 10 cycles of 30 sec ON and 30 sec OFF for NPs. Correct DNA fragments enrichment at around 200bp was verified using the Agilent 2200 Tape Station system (Agilent, G2964AA, 5067-5584 and 5067-5585) and by gel electrophoresis. Three independent biological replicates were performed for each experiment. Antibody references: CTCF (Diagenode, C15410210), OCT4 (Diagenode C15410305), SOX2 (Santa Cruz, sc-365823), CREB (Abcam, ab31387), NRF1 (Abcam, ab55744). Primer sequences for qPCR are listed in **supplementary table 3**.

### *In vitro* methylation assay

Complementary single-stranded DNA (ssDNA) oligos, the forward strand containing one methylated CpG dinucleotide within or directly next to the PF motif were synthetized by Microsynth AG. Hemi-methylated double-stranded DNA (dsDNA) probes were then produced by annealing these ssDNA oligomers. Annealed dsDNA probes were quantified using the Qubit^TM^ 3.0 Fluorometer with the Qubit^TM^ dsDNA HS Assay Kit (ThermoFischer Scientific, Q32854) and diluted to a final concentration of 800nM. Reaction buffer was prepared as follows: 3 Ci/mmol SAM[3H] (Perkin Elmer, NET 155V250UC), 1x methylation buffer [40mM Tris/HCl pH 7.5 (Invitrogen, 15504020), 10mM EDTA (Applichem, A1104-0500), 10mM DTT (Applichem, A1101.0005), 0.2% Glycerol (Sigma, 49767-1L)], 0.2mg/mL BSA (Applichem, A1391,0500), 1x Protease Inhibitor cocktail (ROCHE, 05056489001). 16.68 pmoles of dsDNA probe were added to the buffer in three conditions: 1) buffer only; 2) buffer + DNMT1; 3) buffer + DNMT1 + 1x TF protein at an equimolar concentration to the probe. Samples were incubated at 37°C for 1h, then purified by phenol (Invitrogen 15513-039) and chloroform:IAA (Sigma, C0549-1PT) and ethanol precipitation. The DNA pellets were resuspended in 20μL of TE buffer, then 15μL of the eluate were placed on a filter paper and air dried, the remaining eluate was used to quantify the probe concentration for normalization. The filter papers were transferred into Scintillation Vials (Sigma, V6755-1000EA) with 4.5ml of Ultima Gold Scintillation Liquid (Perkin Elmer, 6013151). Incorporation of ^3^H was measured on a Liquid Scintillation Counter (Wallac, 1409) for 5min. The resulting measurements were normalized to the concentration of the eluate before further normalization to the baseline activity measured in the second condition containing only DNMT1. Recombinant proteins used in the assay were: DNMT1 (Abcam, ab198140), KAISO (Abcam, ab160762), ERα (Abcam, ab82606), NFYA (Abcam, ab131777), E2F1 (Abcam, ab82207), OCT4 (Abcam, ab169842), SOX2 (Abcam, ab169843), NRF1 (Abcam, ab132404), CTCF (Abcam, ab153114), FOXA1 (Abcam, ab98301), SOX9 (Abcam, ab131911), FOXD3 (Abcam, ab134848), KLF4 (Abcam, ab169841), ETS1 (Abcam, ab114322), KLF7 (Abcam, ab132999), NANOG (Abcam, ab134886), OTX2 (Abcam, ab200294), SOX17 (LSBio, LS-G69322-20), CREB (LSBio, LS-G28015-2), GATA3 (LSBio, LS-G67133-20).

### Protein production

Recombinant proteins used in *in vitro* replication experiments were either purchased (SOX2, Abcam, ab169843) or prepared in Sf9 cells. Baculoviruses for expression of Flag-NFYA, Flag-FoxA1, or Flag-CTCF were used to infect 1L of Sf9 cells for 48-72 hours as previously described^71^. Cells were collected by centrifugation and washed with 5-10 volumes of PBS + 0.1mM PMSF. Cells were spun down, washed once with 1X PBS, and pellets were resuspended in 2-3 volumes of Buffer F (20mM Tris pH

8.0, 500mM NaCl, 4mM MgCl_2_, 0.4mM EDTA, 20% glycerol) plus NP40 to 0.05% with protease inhibitors (0.2mM PMSF, 13.5μM TLCK, 0.1μM Benzamidine, 3μM Pepstatin, 55μM Phenanthroline, 1.5μM Aprotinin and 23μM Leupeptin), ZnCl_2_ (10μM final concentration) and DTT (1mM final concentration). Cells were incubated on ice for 30 min and homogenized with a total of 3*10 strokes during the incubation. Extracts were centrifuged (30 min 48000 g), flash frozen, and stored at −80°C. For anti-FLAG affinity purification, extracts incubated with protease inhibitors and 1-2ml of packed anti-FLAG resin (M2-agarose, Sigma), then binding was allowed to proceed overnight at 4°C with rotation. Beads were centrifuged (1500*g for 5 min), then washed with the following series: 2x Buffer FN, 2x BC1200N, 2x BC2000N (the second wash incubated for 15 min), 1x BC1200N, 1x BC600N, 1x BC300N, 1x BC300. The initial washes were carried out in batch (5 min rotation followed by centrifugation at 1000*g for 4 min), and beads transferred to an Econo Column (Bio-Rad) at the BC2000N step. Proteins were eluted by incubation overnight with 0.4mg/mL FLAG peptide in BC300 with 10μM ZnCl_2_ and protease inhibitors. Two additional elutions (with 1h incubations) were collected. Eluted proteins were concentrated using Amicon Ultra 0.5 Centrifugal filter units (10kDa MWCO) (Millipore) and NP40 was added to 0.05% before aliquoting, flash freezing, and storing at −80°C. Protein concentration was determined by Bradford assay, and adjusted for the purity as determined on SYPRO Ruby stained SDS-PAGE gels.

### *In vitro* DNA replication assay

For large scale DNA replication reactions used for bisulfite sequencing, TFs were pre-bound to 100-200ng plasmid template in 60mM KCl, 12mM Hepes pH 7.9, 2mM MgCl_2_, 1mM DTT, 0.12mM EDTA, 12% glycerol, 0.01% NP40, and 10ng/μL DNA template for 15 min at 30°C. For EMSA, 0.5μL of each reaction (5ng DNA) were removed, mixed with 4μL of 50% glycerol/10mM EDTA, and loaded on a 0.8% agarose (SeaKem)/0.5 X TBE gel, which was run for 90 min at 50V. Replication mix was added to the remainder of the reaction. Replication mix consists of (per 100ng DNA): 10μL HeLa S240 extract, 1.38μL replication cocktail (200μM each rNTP, 100μM dATP, dGTP, dCTP, 20μM dTTP, 40mM phospho-creatine, 1ng/μL creatine kinase (Sigma), 3mM ATP, 5mM MgCl_2_), 0.2μL human Topoisomerase II (TopoGen), 1mM DTT, 0.32μL Biotin-18-dUTP or Biotin-11-dUTP (1mM, Jena Bioscience or Fisher), SAM[3H] (1μCi/100ng DNA, Perkin Elmer). Replication reactions were incubated 90 min at 37°C. Replication reactions were stopped with DSB-PK (5μg/µL of proteinase K (Biobasic), 1% SDS, 50mM Tris-HCl pH 8.0, 25% glycerol and 100mM EDTA), digested overnight at 50°C, followed by at least 30 min with RNaseA (1μg/100ng DNA) at 37°C, and purified by phenol-chloroform and chloroform extraction, followed by ethanol precipitation.

For binding to monovalent streptavidin beads (BcMag Monomeric Avidin Magnetic Beads, Bioclone Inc.), 40μl of beads/750 ng reaction were prepared according to the manufacturer’s instructions. Briefly, beads were washed 1X with 4 volumes of ddH_2_0 and 1X with 4 volumes of PBS. All wash and binding steps were carried out at room temperature. Beads were incubated with 3 volumes of 5mM Biotin (in TE-100), followed by washing with 6 volumes of 0.1M Glycine pH 2.8. Beads were then washed twice with 4 volumes of TE-1000mM NaCl and added to purified DNA samples. One sample volume of TE-1000 was added to increase the [NaCl] to facilitate binding. Binding was carried out for at least one hour, and up to overnight with continuous rotation. Beads were washed three times with TE-100 and eluted 3 times with 75μL mM Biotin in TE-100. Elutions were incubated at 50°C with vortexing every 10-15 min.

To measure the incorporation of radioactive SAM[3H] during replication, reactions were carried out as above in the presence of SAM[3H] with 100ng of template per reaction; all steps were scaled down linearly. SAM[3H] incorporation was measured by scintillation counting, and an aliquot of the purified DNA quantified from agarose gels.

### RNA extraction and cDNA preparation

RNA was extracted using the Qiagen RNeasy mini kit (Qiagen, 74104) with the addition of the DNase step (RNase-free DNase set, Qiagen, 79254). RNA integrity was verified by running an aliquot on a 1% agarose gel. Conversion of 1μg of RNA to cDNA was done using the Takara PrimeScript 1^st^ strand cDNA synthesis kit (Takara, 6110A) according to manufacturer’s protocol. qRT-PCR was performed using the StepOnePlus qPCR by Applied Biosystems (ThermoFisher, 4376357) with the Applied Biosystems SYBR^TM^ Green PCR Mastermix (ThermoFisher, 4309155). Primer sequences for qPCR are listed in **supplementary table 3**. Sequencing libraries were prepared from 500ng of RNA using the TruSeq mRNA stranded kit (Illumina, RS-122-2101). Molarity and quality were assessed by Qubit and Tape Station. Biological replicates were barcoded and pooled at 2nM and sequenced on 2 lanes using the Illumina HiSeq 4000 sequencer.

### Motif design

Criteria for choosing WT TF motifs were the following: 1) when available, motifs identified from ChIP-Seq data were selected. 2) if no such data is available, Position Frequency Matrices (PFMs) were obtained from the JASPAR Core Vertebrate 2016 database ^18^; alternatively, from the UniProbe ^19^ or TRANSFAC ^20^ databases. WT motifs were chosen mainly as the consensus sequence found in JASPAR database. To minimize the cross-matching between motifs, we checked that the WT-core motifs (*e.g*. GAATGTTTGTTT) and the combination restriction site-motif-barcode (*e.g*. catgtaGCATGCtgagaaGAATGTTTGTTTtgagaaGCTAGCcatgta) did not match with JASPAR motifs other than intended. This was done using the countPWM() function of the R Biostrings package using min.score=”90%”. Scrambled (Sc) motifs were created by random shuffling of the WT motif except for the CG dinucleotides. The number and position of CG dinucleotides were maintained in WT and Sc motifs. For example, WT: C**CG**TAGTCGA and SC: T**CG**AGCAGTC. Score and SC motif-barcode combination were also checked for cross-matching with other JASPAR motifs as for WT-sequences. To ascertain how closely the WT or Sc sequences match with the respective motif’s PFM, a “normalized score” was defined. At each position in WT or Sc sequence, the probability of corresponding nucleotide in the PFM was taken as the match score for that position. The average of match scores for all positions was defined as “normalized score”. Normalized scores of WT-sequences were high (>0.7) and only those Sc motifs whose normalized score were at least 0.3 lower than the corresponding WT motif, were used.

### Library Data Processing

Paired-end libraries were trimmed using Trim Galore^72, 73^ and reads with a quality score below 20 were discarded. Demultiplexing was performed with Flexbar^74, 75^, using the 6bp library barcode plus 4bp of the neighbouring adapter for identification, with 0 mismatches allowed. Concomitantly, reads were tagged using to the UMI-tags option of Flexbar based on the 8 Unique Molecular Identifier (UMI) nucleotides that follow the library barcode sequence. Prior to mapping, motifs were extracted and PE reads were classified according to their motif using the *vmatchPattern* function with unfixed sequences (allowing IUPAC code for CpGs inside the motifs) from the BioString^76^ package designed for R^77^, with 0 mismatches allowed. Reads were then mapped using Bowtie2^78^ and Bismark^79^, filtering out reads with non-CG methylation below 2%. This filtering step was not performed for the libraries generated in TET TKO cells as the levels of non-CG methylation was significantly higher in these cells (**supplementary figure 3b**). Reads were then deduplicated based on their UMI tag using the UMI_tools software^80^ to remove PCR amplification biases. The percentage of methylation for each CpG position was extracted using Bismark, considering a minimum coverage of 10 reads (**supplementary figure 2b**). Biological and technical replicates were pooled to ensure sufficient coverage upon verification by Multi-Dimensional Scaling (MDS) that replicates were clustering well together^81^. Ascending hierarchical clustering of the motifs, based on the methylation data, was obtained using the *hclust* function in R.

### RNA-Seq analysis

SE 50bp reads were trimmed^82^ and then mapped to the mouse reference genome (GRm38.89 version from Ensembl) using the RNAseq aligner STAR^83^ and featureCounts^84^ to assign reads to their genomic features. Library size normalization and calculation of differential gene expression were performed using the edgeR package. Genes with a normalized maximal expression of less than 1 RPKM in all replicates were discarded. Fold-change and Benjamini-Hochberg corrected p-value thresholds were set respectively to 3 and 1‰ for the differently expressed genes.

### Statistical analysis

For the NGS data, methylation differences between WT and Sc for each CpG and for each motif were calculated using the DSS (Dispersion shrinkage for sequencing data) package^85, 86^, with thresholds for Δmeth and corrected p-value fixed respectively at 10% and 5%. The percentage of methylation of each CpG was smoothed with adjacent CpG to improve mean estimation. The smoothing option was applied to a range of 50bp. Hypomethylated regions in WT vs Sc conditions (HMRs) were defined as regions of more than 50bp and containing a minimum of 3 consecutive CpGs, each having a Δmet (%met_WT – %met_Sc) of 10% or higher.

For *in vitro* methylation assays, three biological replicates were analysed using two-tailed unpaired t-test with Welch correction. *In vitro* replication results were analysed by one-sample unpaired t-test.

## Supporting information

SupFig1

SupFig2

SupFig3

SupFig4

SupFig5

SupFig6

## Acknowledgements

We thank M. Lorincz (University of British Columbia-Vancouver) for providing the RMCE plasmids; Ann Dean (National Institute for Diabetes and Digestive and Kidney Diseases) and D. Schübeler (Friedrich Miescher Institute) for the RMCE target ES cell line; R. Jaenisch (Whitehead Institute for Biomedical Research) for providing the TET-TKO ES cells. We are grateful to Sylvain Lemeille for help in designing the experimental approach and to Z. Herceg, T. Baubec, G. Andrey, and S. Braun for helpful comments on the manuscript. Sequencing was performed at the iGE3 Genomic Platform of the University of Geneva. LV was supported by the iGE3.

Research in the laboratory of RM is funded by the SNSF grants PP00P3_150712, PP00P3_179063 and PP00P3_190075, the Boninchi Foundation, Von Meissner Foundation, and the Novartis Foundation for Medical-Biological research.

## Author Contributions

RM and LV conceived the study; LV, HS, NF and RM performed experiments and analysed the results; HS and RM performed the in vitro methylation assays and contributed to the other experiments; NF performed the in vitro replication assays; VY performed the bioinformatic analysis; SA designed the motifs and contributed to the bioinformatic analysis; LV inserted the RMCE selection cassette into the TET-TKO ES cell lines; LV, VY and RM prepared the figures and wrote the manuscript with input from all authors.

---

**Supplementary Figure 1. Experimental approach. (**a**)** Schematic representation of the Hi-TransMet approach. A library of targeting plasmids each containing the same bacterial DNA (grey line) but a different motif (colored rectangles) and barcode (dashed box) and flanked by *LoxP* sites (triangles) were transfected into ESCs containing the RMCE site together with a plasmid expressing CRE recombinase. This leads to the replacement of the selection cassette (Hygromycin/Thymidine Kinase) by the bacterial fragment. Ganciclovir treatment selects the cells that underwent recombination. Genomic DNA is extracted from successfully-recombined cells and treated with sodium bisulfite. (**b**) Sequence of the bacterial fragment FR1 used in this study. Primer sequences for Bisulfite PCR and library preparation are indicated in green. US_Primer is the upstream primer pair, DS_Primer is the downstream primer pair (please refer to text for further details). Edits to the original sequence^42^ are indicated in dark blue (additions) and light blue (changes of position). The fragment was inserted into the RMCE donor plasmid by directional cloning using the restriction enzymes BamHI and HindIII (flags); motifs were later inserted by directional cloning with SphI and NheI (flags).

**Supplementary Figure 2. Hi-TransMet library preparation and molecular barcoding.** (**a**) NGS library preparation. Step 1, UMI assignment. A target-specific reverse primer, including of a UMI tag (colored Ns) and a library barcode (black) is annealed to the bisulfite-converted DNA and the target sequence is extended. Step 2, non-barcoded amplification. A short PCR amplification is performed using forward target-specific primers and a reverse universal primer. Step 3, addition of sequencing adapters by PCR amplification. Unused primers and primer dimers are removed between each step. The region of interest surrounding the motifs is PCR amplified using two sets of universal primers, upstream and downstream of the motifs, covering about 500 bp flanking the binding sites. (**b**) Overview of the bioinformatic pipeline.

**Supplementary Figure 3. Overall methylation levels on FR1 fragment in WT ESCs, NPs and TET TKO ESCs** (**a**) Mean ratio of CpG methylation in WT ESCs, NPs and TET TKO ESCs measured at FR1 fragments containing Sc motifs. P-values (two-tailed paired t-test): NPs/-SssI vs ESCs/-SssI: p<0.0001; NPs/+SssI vs ESCs/+SssI: p=0.0025; TET-TKO/+SssI vs ESCs/+SssI: p<0.0001; TET-TKO/+SssI vs NPs/+SssI: p=0.1527. (**b**) Mean ratio of non CpG methylation (CHG+CHH) in WT ESCs, NPs and TET TKO ESCs measured at FR1 fragments containing Sc motifs. P-values (two-tailed paired t-test): NPs/-SssI vs ESCs/-SssI: p=0.0010; NPs/+SssI vs ESCs/+SssI: p<0.0001; TET-TKO/+SssI vs ESCs/+SssI: p=0.0002; TET-TKO/+SssI vs NPs/+SssI: p<0.0001.

**Supplementary Figure 4. Differentiation into neuronal progenitors** (**a, b**) Heatmaps representing methylation percentages at WT and Sc motifs in the FR1/-SssI condition (**a**) and FR1/+SssI (**b**). (**c**) qPCR validation of NP differentiation by analysis of ESC-specific (Oct4, Nanog and Sox2) and NP-specific markers (FoxA1, Pax6 and Sox9) (biological triplicate).

**Supplementary Figure 5. *In vitro* replication assay** (**a**) Schematic representation of the *in vitro* replication assay. A bacterial DNA fragment is cloned into the SV40 replication vector containing an origin of replication. The resulting plasmid is *in vitro* methylated using the SssI methyltransferase. Plasmid is then incubated with the SPF of interest, then replication occurs in the presence of T-Antigen and cell extract. Biotinylated, replicated DNA is purified by immunoprecipitation using streptavidin beads and DpnI digestion of the non-replicated plasmid (**b**). (**c**) SPF binding to the plasmid was verified by EMSA. (**d**) DNA methylation is maintained during *in vitro* DNA replication. –SssI and +SssI plasmids were incubated in HeLa extracts in the presence of SAM[3H] and in the presence or absence of T-Ag. DNA was purified and SAM[3H] measured by scintillation counting. Graph shows CPM - background. (**e**) Bisulfite Sanger sequencing analysis of the region surrounding the binding sites in absence or presence of replication (+/-T-Ag) and the PF (+/-PF). Vertical bars correspond to CpG positions, and the color code corresponds to the percentage of methylation calculated for each CpG with a minimum coverage of 10 bisulfite reads.

**Supplementary Figure 6. ChIP validation of PF binding** (**a**) Chromatin immunoprecipitation (ChIP) to validate CREB binding in ESCs and NPs. Single-motif containing ESCs were generated by RMCE insertion of FR1/CREB_WT and FR1/CREB_Sc fragments individually. Cells were then differentiated into NPs. Results are shown as mean + SEM of three biological replicates. P-values (two-tailed unpaired t-test): WT_NPs/-SssI vs Sc_NPs/-SssI: p=0.0202; WT_ESCs/-SssI vs WT_NPs/-SssI: p=0.0168; WT_ESCs/+SssI vs WT_NPs/+SssI: p=0.0340. (**b**) SOX2 ChIP in ESCs and NPs containing the OCT4-SOX2 motif. P-value WT_ESCs/+SssI vs Sc_ESCs/+SssI: 0.0012 (two-tailed unpaired t-test) (**c**) OCT4 ChIP in ESCs and NPs containing the OCT4-SOX2 motif. (**d**) SOX2 ChIP in SOX2-motif containing ESCs. (**c**) NRF1 ChIP in NRF1-motif containing ESCs.

**Supplementary Table 1.**
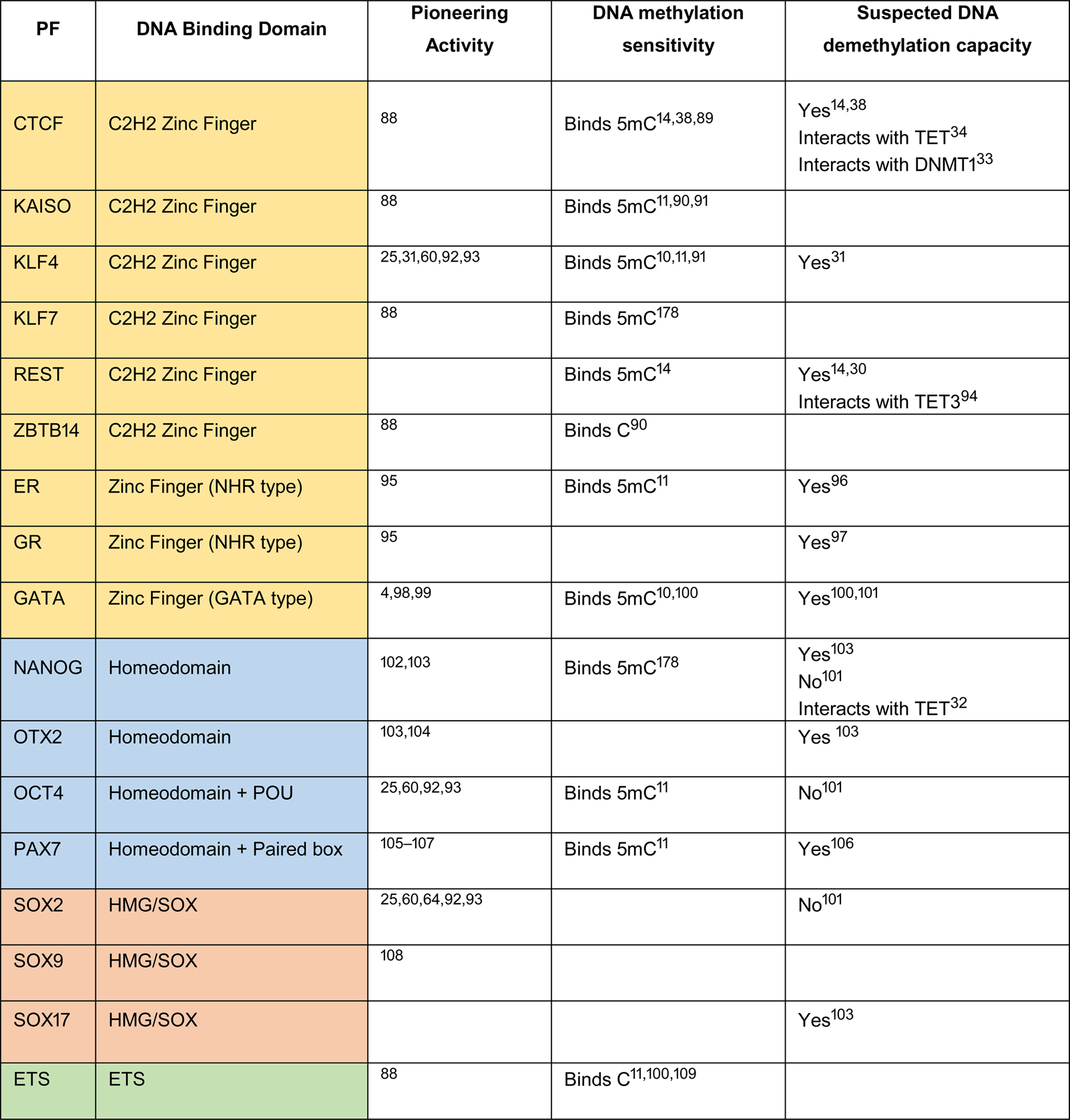
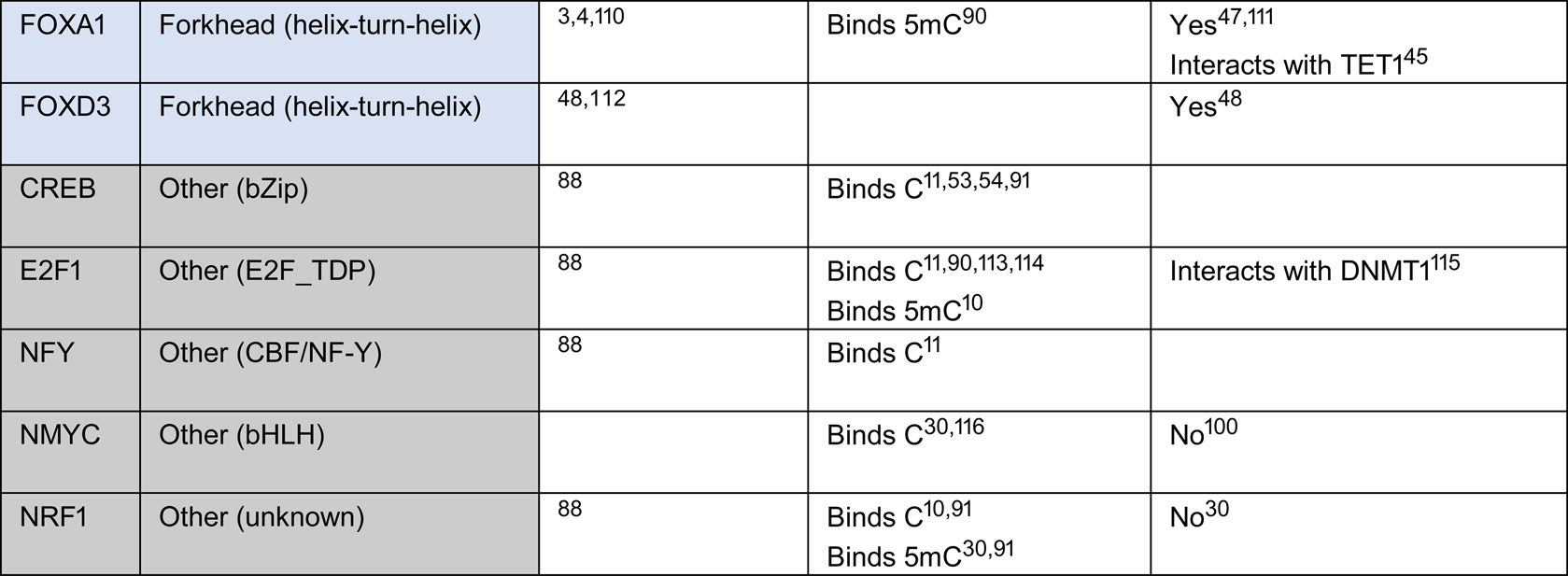
**List of PFs selected for RMCE screening**. DNA binding domains are reported as indicated in the Human TFs database (http://humantfs.ccbr.utoronto.ca/)^87^. For each factor, the main references for: pioneering activity; sensitivity or binding to methylated DNA; suspected DNA demethylation capacity and/or suspected or proven interactions with TETs and DNMTs are indicated

**Supplementary Table 2.**
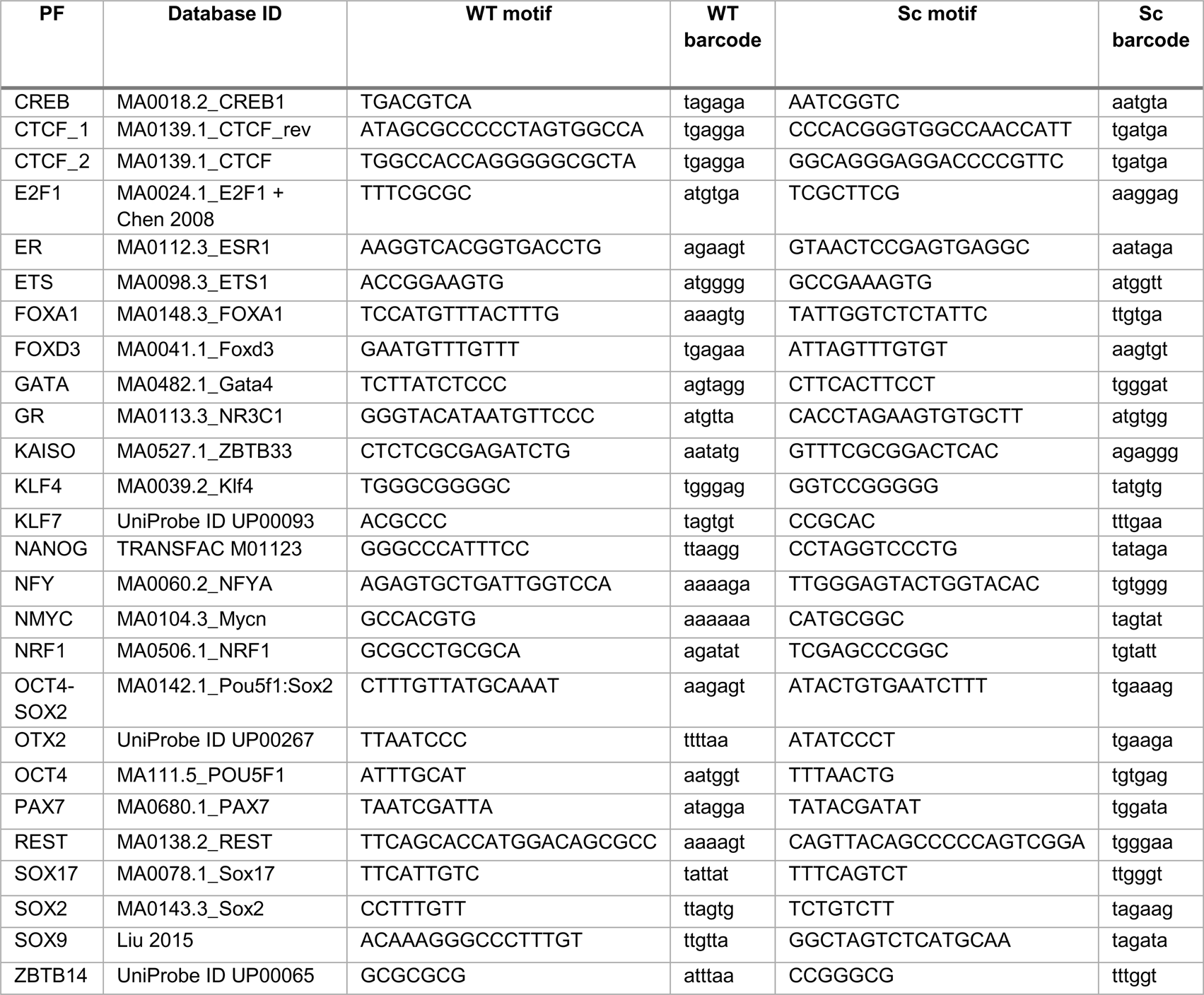
List of WT and Sc PF motifs and barcodes.

**Supplementary Table 3.**
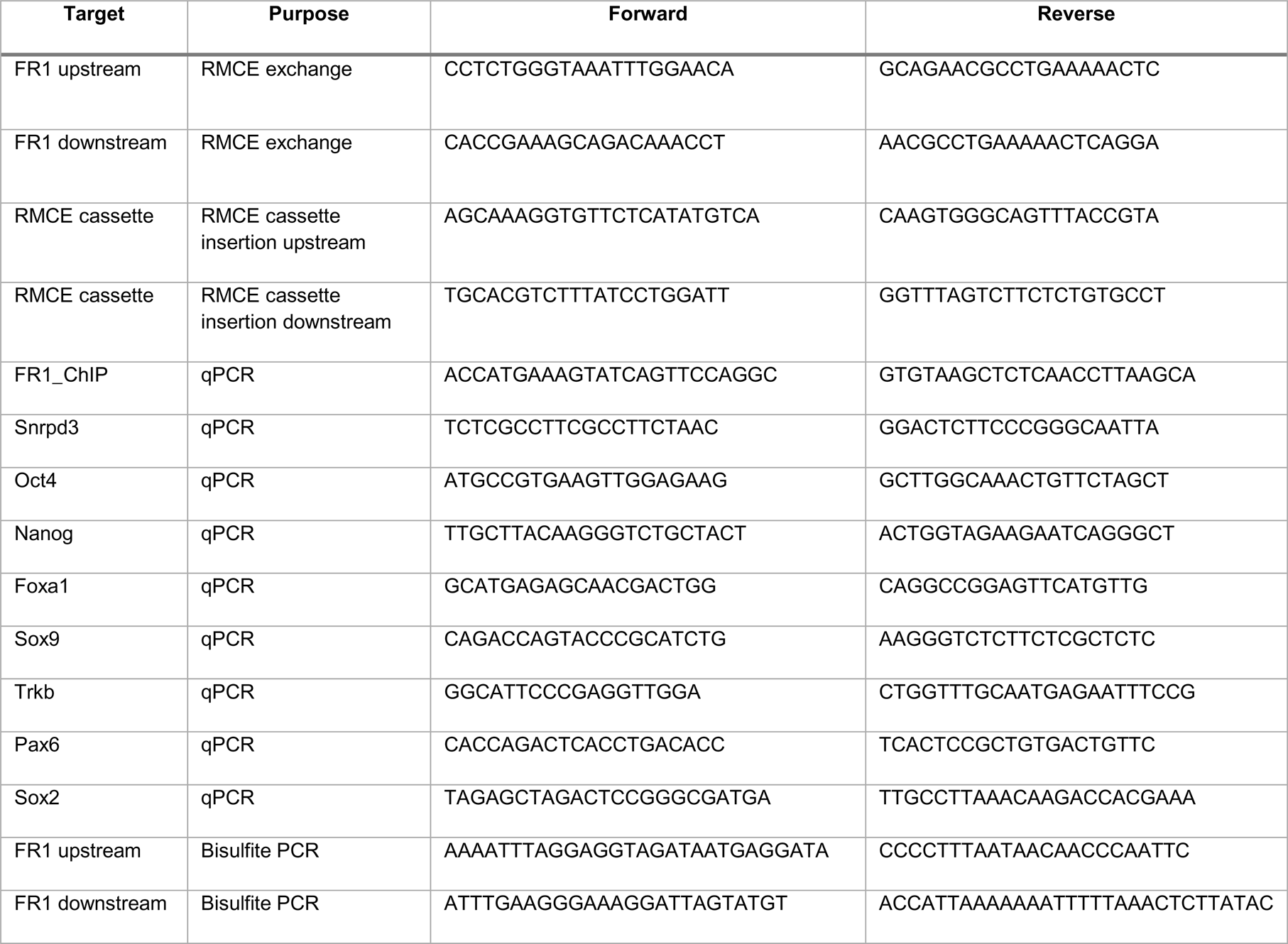
List of primers used in this study.

**Supplementary Table 4.**
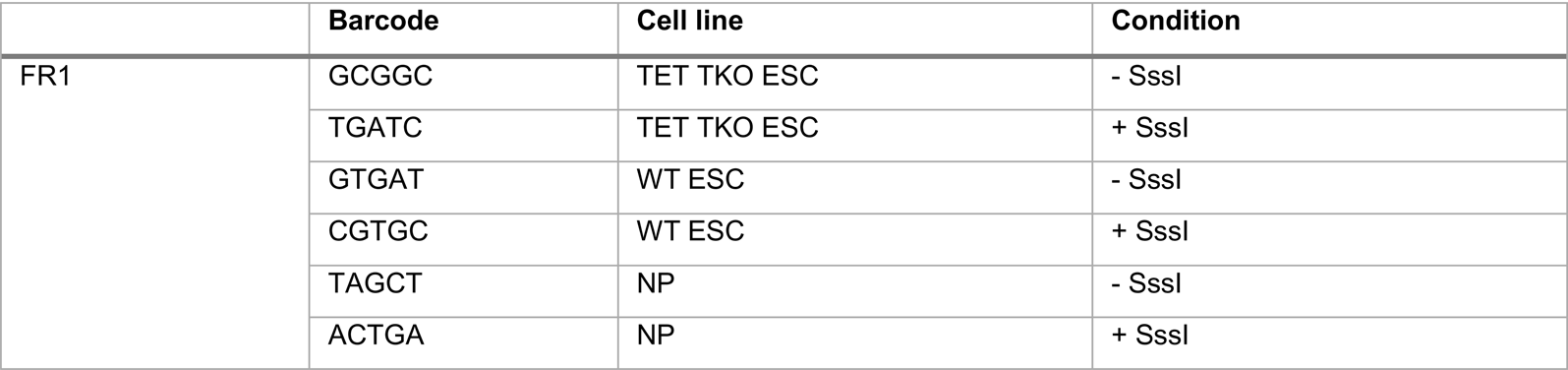
List of library barcodes.

|  | Barcode | Cell line | Condition |
| --- | --- | --- | --- |
| FR1 | GCGGC | TET TKO ESC | - Sssl |
|  | TGATC | TET TKO ESC | + Sssl |
|  | GTGAT | WT ESC | - Sssl |
|  | CGTGC | WT ESC | + Sssl |
|  | TAGCT | NP | - Sssl |
|  | ACTGA | NP | + Sssl |

**Supplementary Table 5.**
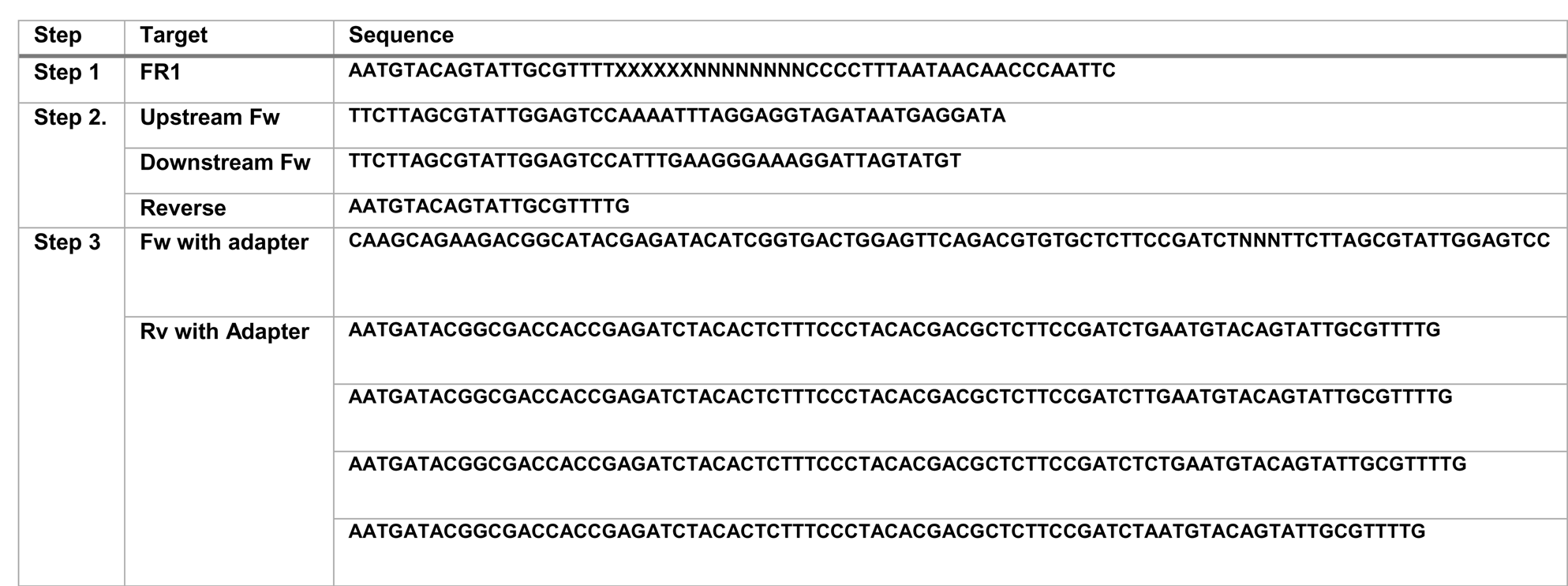
Hi-TransMet library preparation primers.

