## Supplementary figures and images for "High throughput screening identifies SOX2 as a Super Pioneer Factor that inhibits DNA methylation maintenance at its binding sites"

### SupFig1

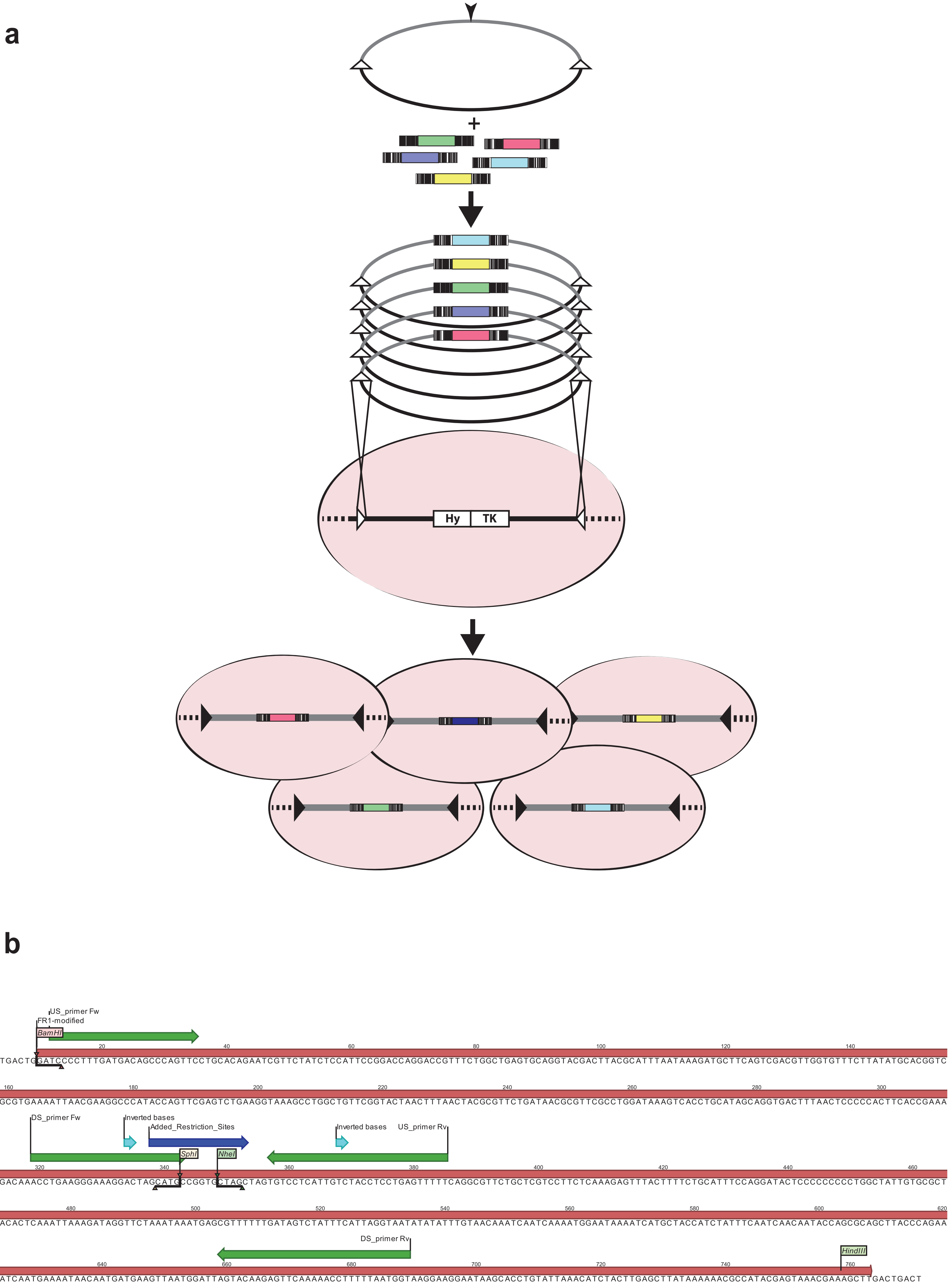


Supp. Figure 1

### SupFig2

a

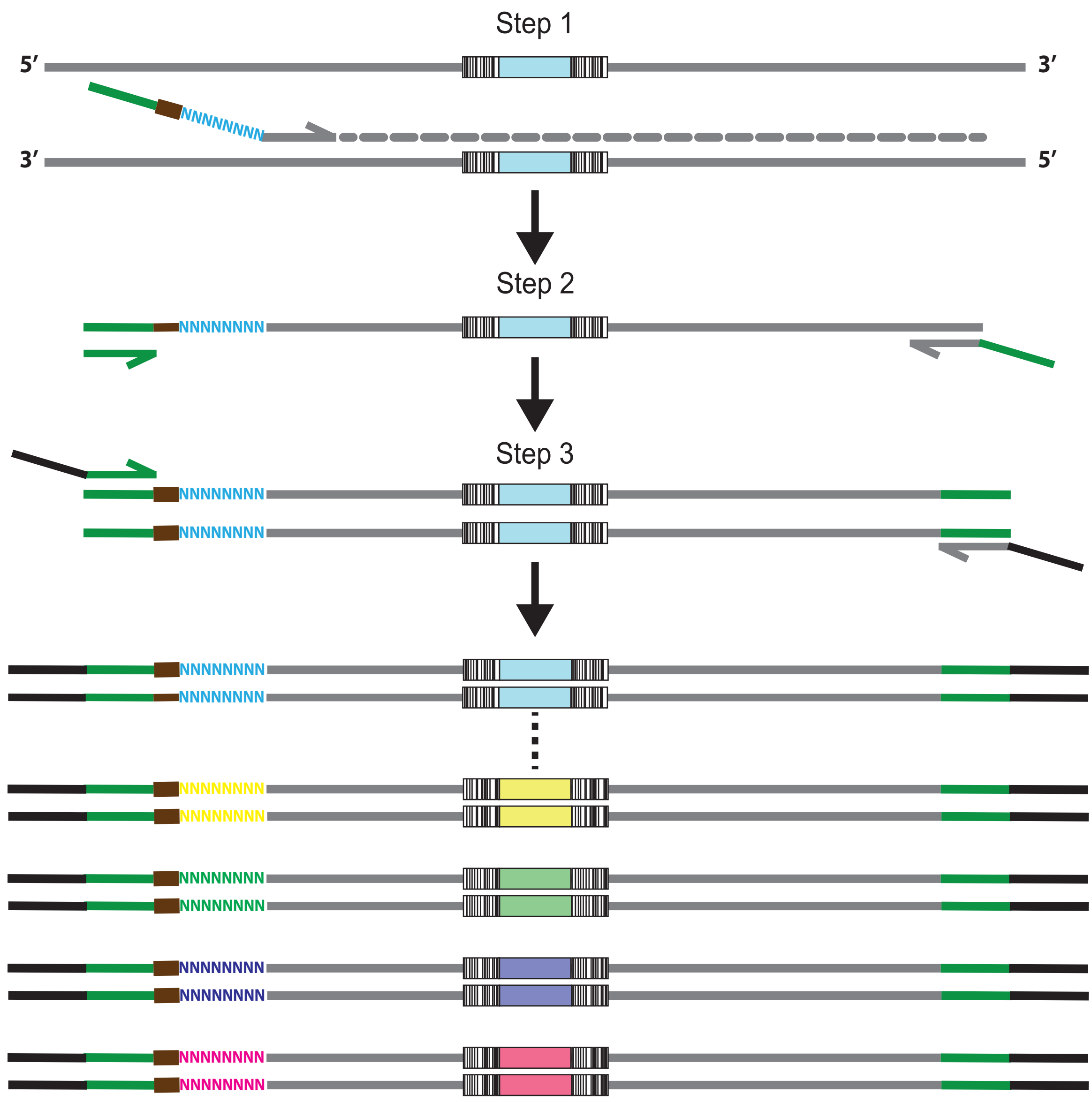


b

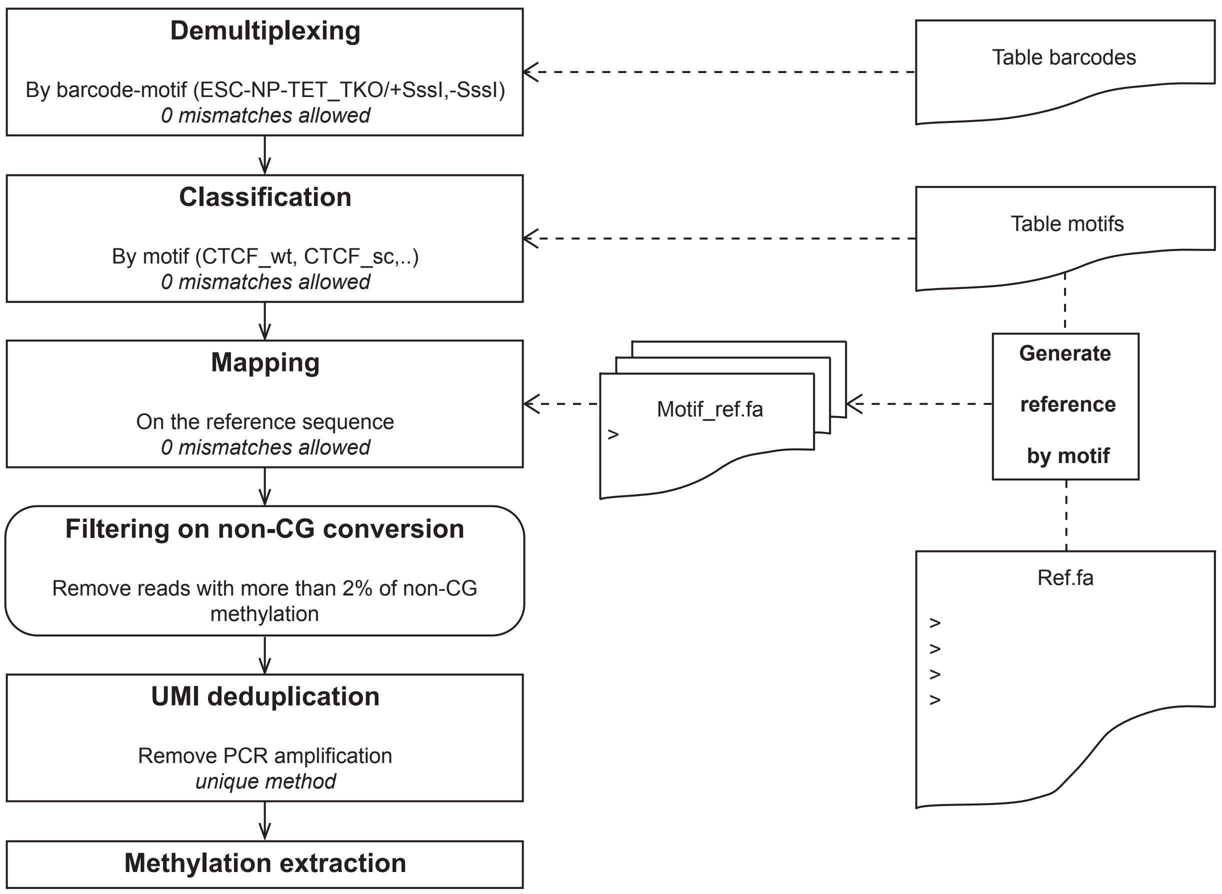


Supp. Figure 2

### SupFig3

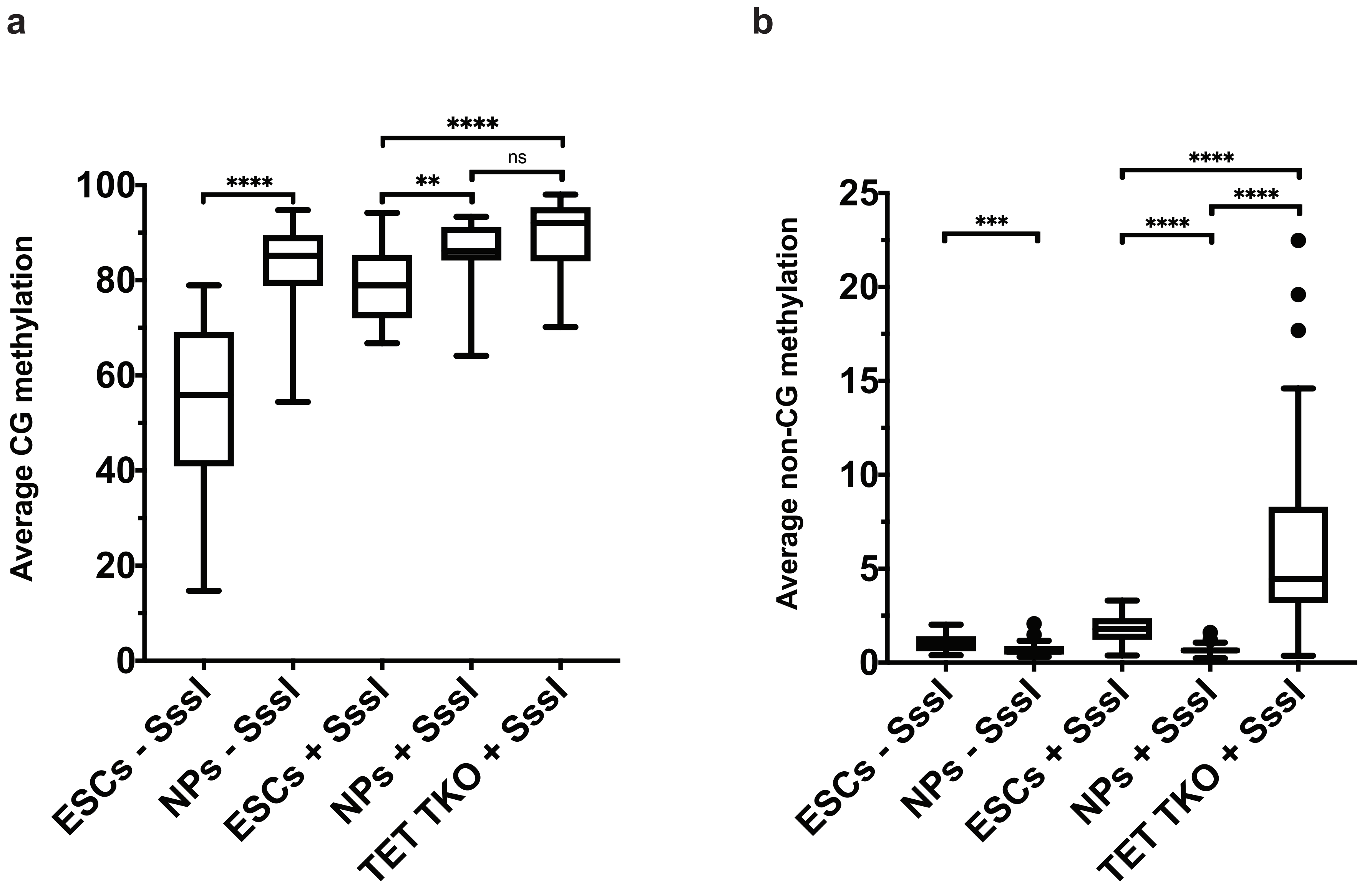


Supp. Figure 3

### SupFig4

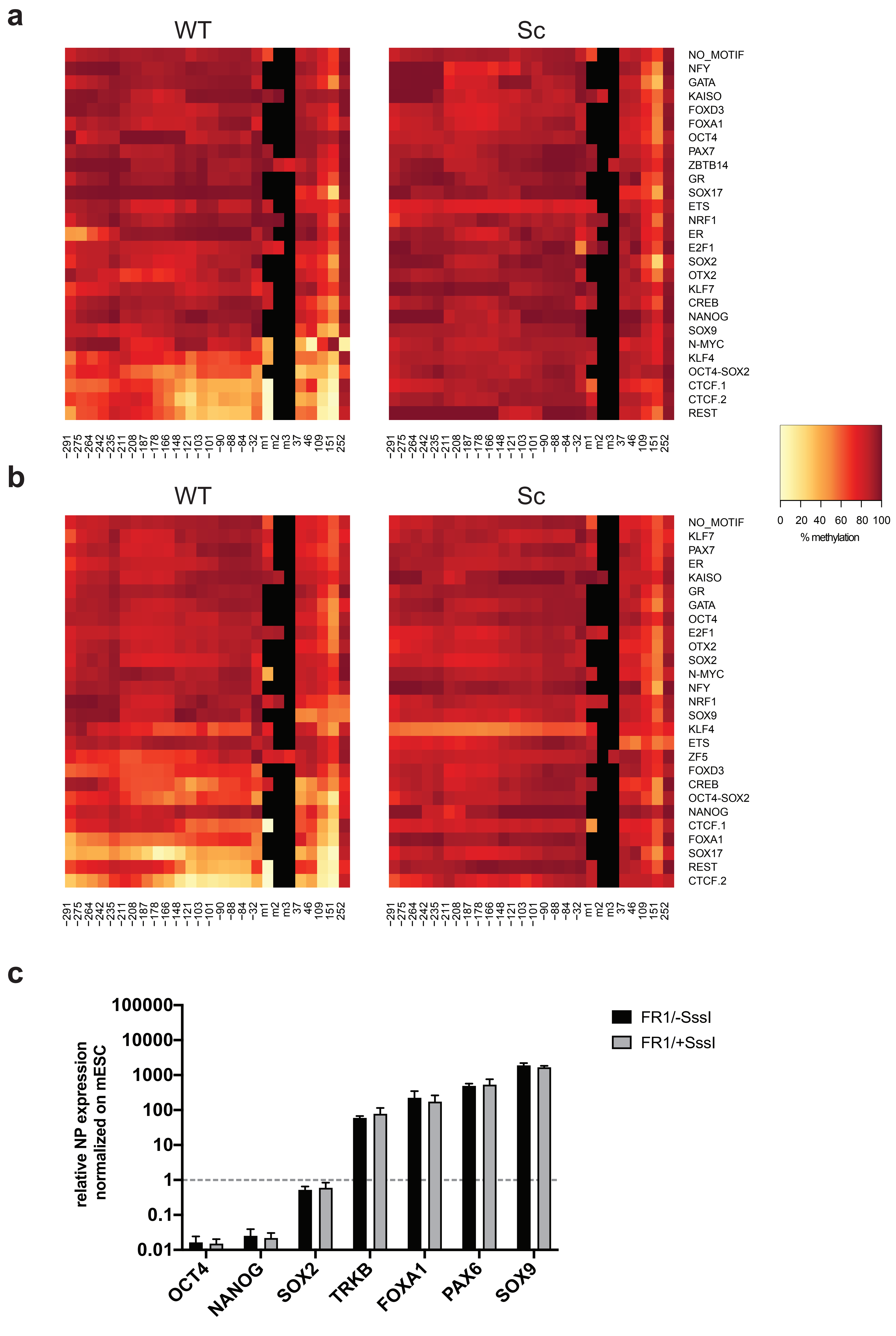


Supp. Figure 4

### SupFig5

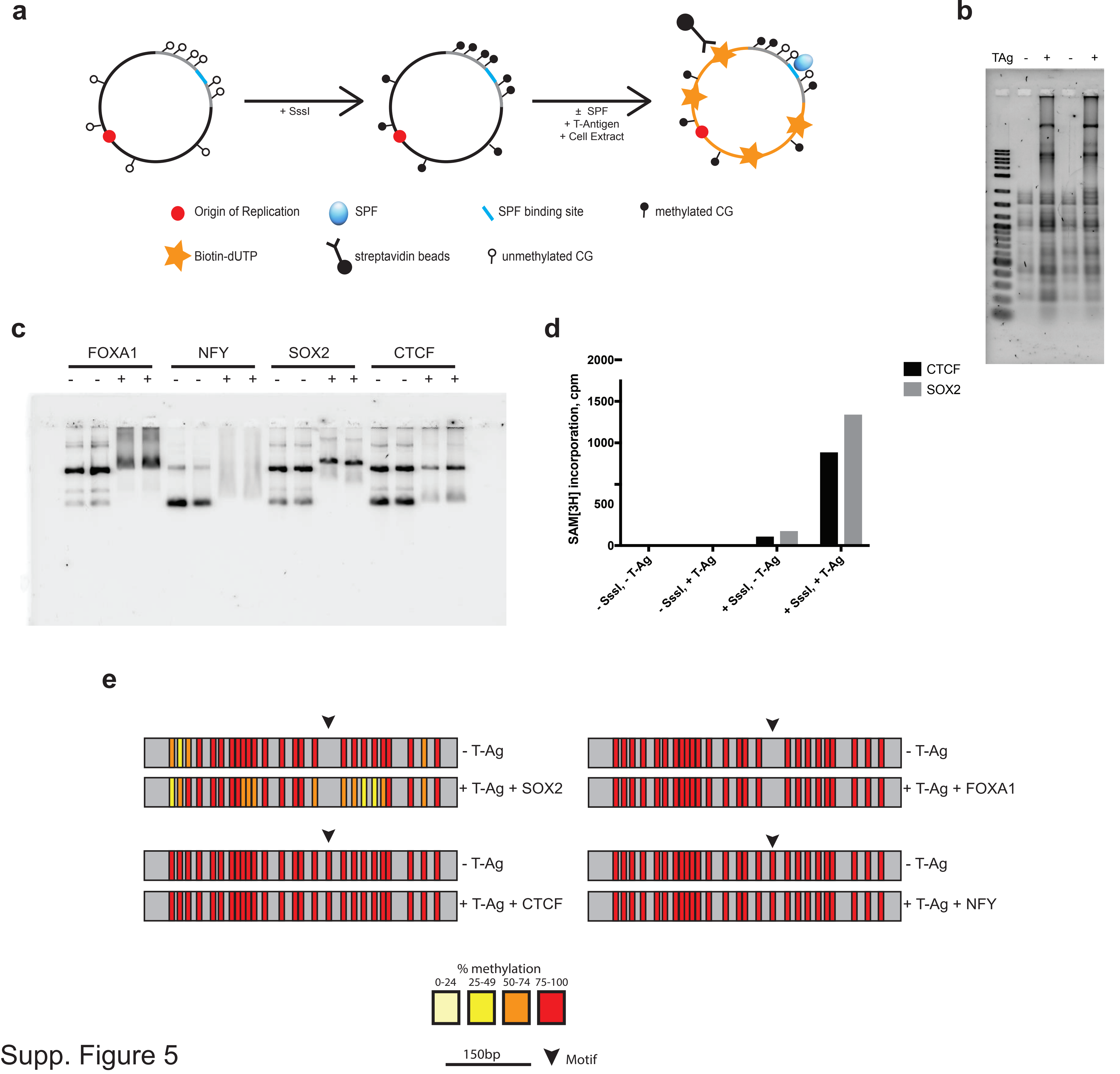


Supp. Figure 5

### SupFig6

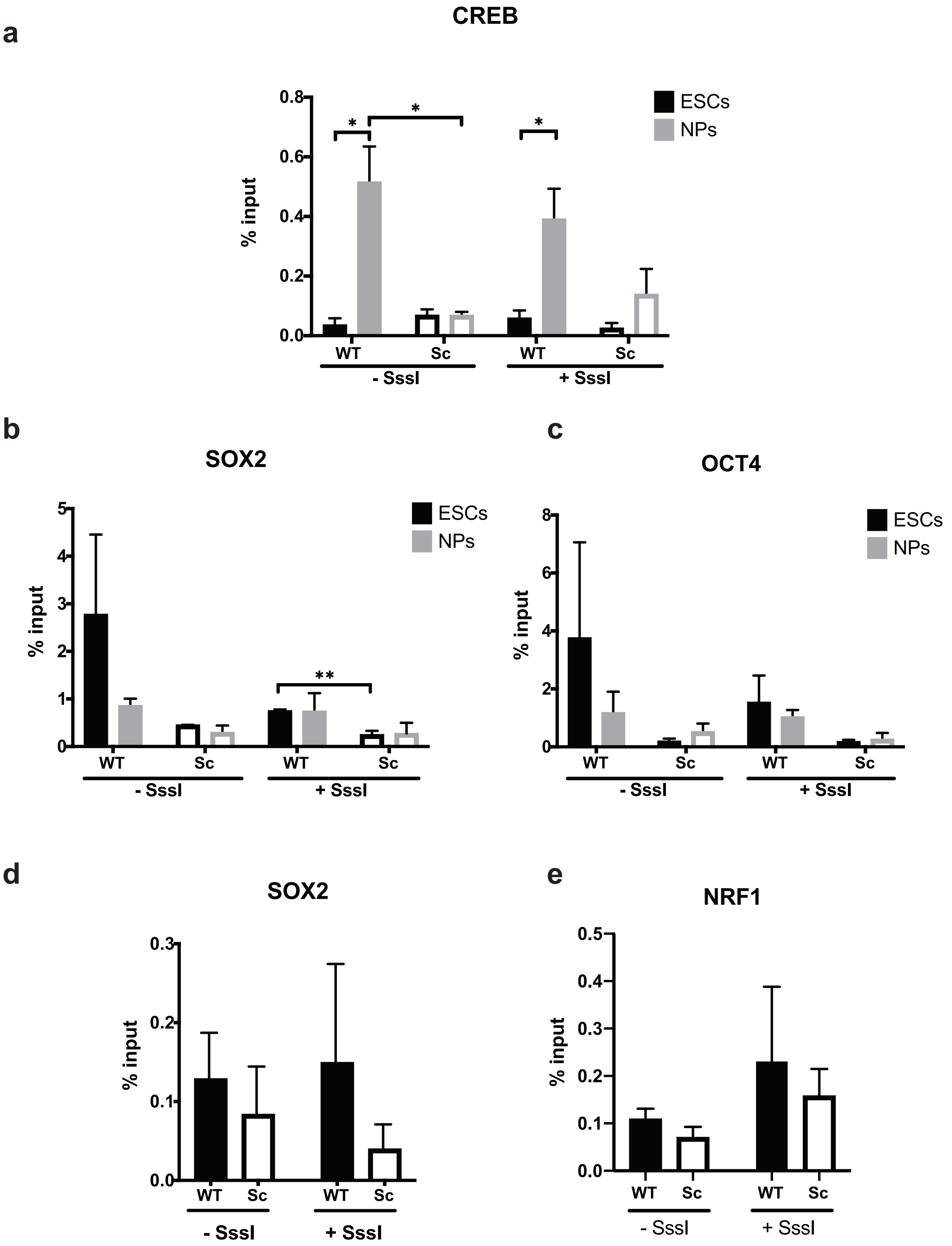


Supp. Figure 6
